# Linking mesozooplankton and SAR11 bacteria in Oxygen Deficient Zones and the open ocean

**DOI:** 10.1101/2022.09.04.506529

**Authors:** Clara A. Fuchsman, Matthew D. Hays, Paulina Huanca-Valenzuela, Benjamin P. Gregory, Louis V. Plough, Megan E. Duffy, Richard G. Keil, Xuefeng Peng

**Author notes:** Corresponding author: Clara A. Fuchsman, University of Maryland Center for Environmental Science, Horn Point Laboratory. Cambridge MD 21613 USA. The authors have no competing interests.

## Abstract

The gravitational biological pump is not large enough to account for microbial heterotrophic activity in the mesopelagic ocean. Migrating zooplankton may be a key source of organic matter transport to depth. Here we show signatures of zooplankton in the suspended organic matter at the zooplankton vertical migration depth in the Eastern Tropical North Pacific Oxygen Deficient Zone (ETNP ODZ). We examine the mesozooplankton community in metagenomic depth profiles using the mitochondrial cytochrome c oxidase (COI) gene as a marker in the ETNP and Eastern Tropical South Pacific (ETSP) ODZs and at the oxic Hawaii Ocean Timeseries (HOT). Additionally, eukaryotic transcripts (polyA-selected) were examined for zooplankton in the ETNP. While zooplankton eDNA increased in the ODZ, zooplankton eRNA decreased in the ODZ, similar to previous net-based data, implying that eDNA is better preserved under anoxia. At all stations, Cnidaria, often missed in net-based data, contributed greatly to the zooplankton eDNA/eRNA. SAR11 abundance, determined from the single-copy core gene (*rpoB*), significantly correlated with zooplankton eDNA, with R^2^ values >0.8 at all stations. Strong correlations between SAR11 and zooplankton have not been previously reported, but are logical as SAR11 bacteria consume and zooplankton excrete simple dissolved organic compounds. SAR11 bacteria possessed genes to utilize urea and taurine in the ODZ, both compounds known to be excreted by zooplankton. In ODZs, SAR11 bacteria preferentially used the taurine degradation pathway leading to C and N assimilation, not the pathway for organic S assimilation, probably due to additional sources of organic S in ODZs.

## Introduction

Oxygen Deficient Zones (ODZs) are regions of the ocean that naturally have <10 nM oxygen at mid-depths [1, 2]. The oxygen content of the Pacific Ocean has been decreasing since the 1980s [3] and ODZs are expanding [4]. Repeat measurements indicate that the ETNP ODZ has thickened over the past 40 years, both shoaling and deepening [5]. Without oxygen, the microbial community in an ODZ relies on nitrate as an electron acceptor, sometimes reducing it to N_2_ gas. Ocean ODZs host 30-50% of marine fixed-N loss through N_2_ production [6]. In ODZs, N_2_ production is mediated by free-living autotrophic anammox (anaerobic ammonia-oxidizing) bacteria (NO_2_ ^-^ + NH_4_ ^+^ → N_2_) and particle-attached heterotrophic denitrifiers (Org C + → 2NO_3_-N_2_ + CO_2_) [7–9]. However, nitrate reduction to nitrite is the most common metabolism in ODZs [7]. The key metabolisms in the ODZ rely on organic matter or ammonium produced from organic matter remineralization [10].

Settling fluxes of organic carbon to the mesopelagic underestimate microbial production in the dark ocean [11, 12]. Fluxes from zooplankton and gelatinous zooplankton have been suggested as one understudied source of carbon to the mesopelagic that could help fix this imbalance [11–13]. Models that include migrating zooplankton consistently show that this migration increases the biological pump, decreases oxygen concentrations at depth and increases respiration at depth [14–16]. Active transport due to zooplankton migration has been modeled to add fluxes equivalent to 10-40% of the particle flux [15, 17], and sinking jellyfish carcasses have the potential to be an unquantified but important C flux [13]. In the Costa Rica Dome, a high productivity anoxic region adjacent to the ETNP, according to inverse modeling of N isotopes, migrating zooplankton were responsible for 21-45% of organic matter sources to anoxic waters at depth [18]. In the offshore ETNP, organic matter is limiting [19, 20]. Zooplankton could be a source of both ammonium and organic matter to ODZs [21, 22].

The total zooplankton biomass in ODZs is reduced compared to overlying oxic waters [23]. However, zooplankton do regularly migrate into the ODZs [21, 23–25]. Anoxia tolerant species are found in many different zooplankton groups. For example, the organisms found in net tows of the ETNP ODZ include pteropods, polychaetes, lantern fish, euphausiids, and copepods [24, 25]. Different copepod species have very different oxygen requirements and depth profiles, including sometimes copepods in the same genus [24]. This is likely true of other zooplankton types as well. Additionally, nets miss or undercount fragile organisms such as Cnidaria (jellyfish) [26]. The anoxia-tolerant zooplankton that migrate into the deep ODZ daily [21, 24, 25] can provide organic matter to the ODZ microbial community by active transport [22] and/or by dying, degrading, and sinking [27] into the ODZ.

Active transport, where animals swim to depth before defecating and excreting, is known to be an important flux of organic matter in the ocean. At HOT (Hawaii Ocean Time-series), an oxic station in the North Pacific subtropical gyre, active transport by excretion from vertically migrating zooplankton provides dissolved N (ammonium and DON) (14% added to sinking particle fluxes), DOC (18%), and P (phosphate and DOP) (41%) to the microbial community [28]. It has been suggested that migrating zooplankton could excrete ammonia at depth in the ODZ, stimulating N_2_ production by anammox bacteria [21]. Indeed, in the ETNP, the depth of the N_2_ concentration maximum at the offshore ODZ (300m) corresponds exactly with the depth of zooplankton vertical migration [29]. However, partial repression of respiration and ammonia excretion under anoxia has been found in crustaceans [30, 31]. At an ETSP station over the continental slope, after partial repression by anoxia was taken into account, and numbers of euphausiids were corrected for net evasion, euphausiids were calculated to excrete 0.7 nM d^-1^ ammonium [32]. As a comparison, N_2_ production rates offshore at 250-500 m were 2.5-3.0 nM N/d in the ETNP and 1.5-3 nM N/d in the ETSP [19, 33]. Thus, these levels of ammonium excretion could potentially be significant but likely not sufficient to explain N_2_ production. However, a variety of small compounds with reduced N are likely excreted at depth. Dissolved amino acids glycine, alanine, and taurine, as well as nucleosides, and urea all are excreted in abundance by zooplankton in oxic waters [34–37]. Zooplankton are likely contributing particulate organic matter (POM), dissolved organic matter (DOM), and ammonia to the microbial community in the ODZ.

Though often forgotten, gelatinous zooplankton also excrete ammonia, phosphate, dissolved organic C (DOC), dissolved organic N (DON), and dissolved organic phosphorus (DOP) [38] and produce significant amounts of fecal pellets [13]. Unlike crustacean zooplankton and pteropods [37], jellyfish such as ctenophores and schyphomedusae do not excrete urea [39], but they do excrete taurine, glycine, and alanine [35]. The ratio of DOC and DON in ctenophores and schyphomedusae excreta can reach a C:N ratio of 29:1 [38]. Along with excretion and egestion, these gelatinous zooplankton also produce mucus which has a C:N ratio of 25:1 [40]. However, despite this excess in C, jellyfish DOM is actively and quickly respired, though bacterial growth efficiency was only 10% [40].

SAR11 are small heterotrophic bacteria with streamlined genomes [41, 42]. SAR11 in the oxic ocean are heterotrophs that utilize oxygen, but SAR11 bacteria in the ODZ reduce nitrate [43]. SAR11 bacteria are both the most abundant heterotrophic bacteria and the most abundant nitrate reducers in the ETNP ODZ, reaching 60% of the community in the water column [7]. Unsurprisingly, SAR11 ecotypes differ between the oxic ocean and in ODZ with clades 1c and IIa.A predominantly living in ODZs [43, 44]. SAR11 bacteria consume only small dissolved organic compounds and can use a C1 metabolism [45]. Because SAR11 bacteria have small streamlined genomes, they need exogenous sulfur for growth [46]. Taurine is one of several sulfonates, organic S compounds with a C-S bond, that can provide S to SAR11, but taurine is also a source of C, N, and energy [47–50]. Some SAR11 bacteria can also utilize urea as an N and C source [51]. As both urea and taurine are excreted by zooplankton [34, 35, 37, 39], they may constitute a direct link between SAR11 and zooplankton.

Recently, the Census of Marine Zooplankton, a curated database of the mitochondrial cytochrome c oxidase (COI) gene across marine zooplankton, was released [52]. Here we created a phylogenetic tree using reference sequences from the Census of Marine Zooplankton and use phylogenetic read placement of the COI gene to examine zooplankton DNA in-depth profiles of cellular metagenomes from the ETNP and ETSP ODZs and the oxic Hawaii Ocean Timeseries (HOT). We then compare these zooplankton profiles to microbial communities. We report significant correlations between zooplankton and the free-living bacteria SAR11, as well as their implications for ODZ biogeochemistry.

## Methods

This work involves metagenomic, metatranscriptomic, and geochemical data from several ODZ stations in the ETNP and ETSP and HOT (Figure 1; Table S1).

**Figure 1.**
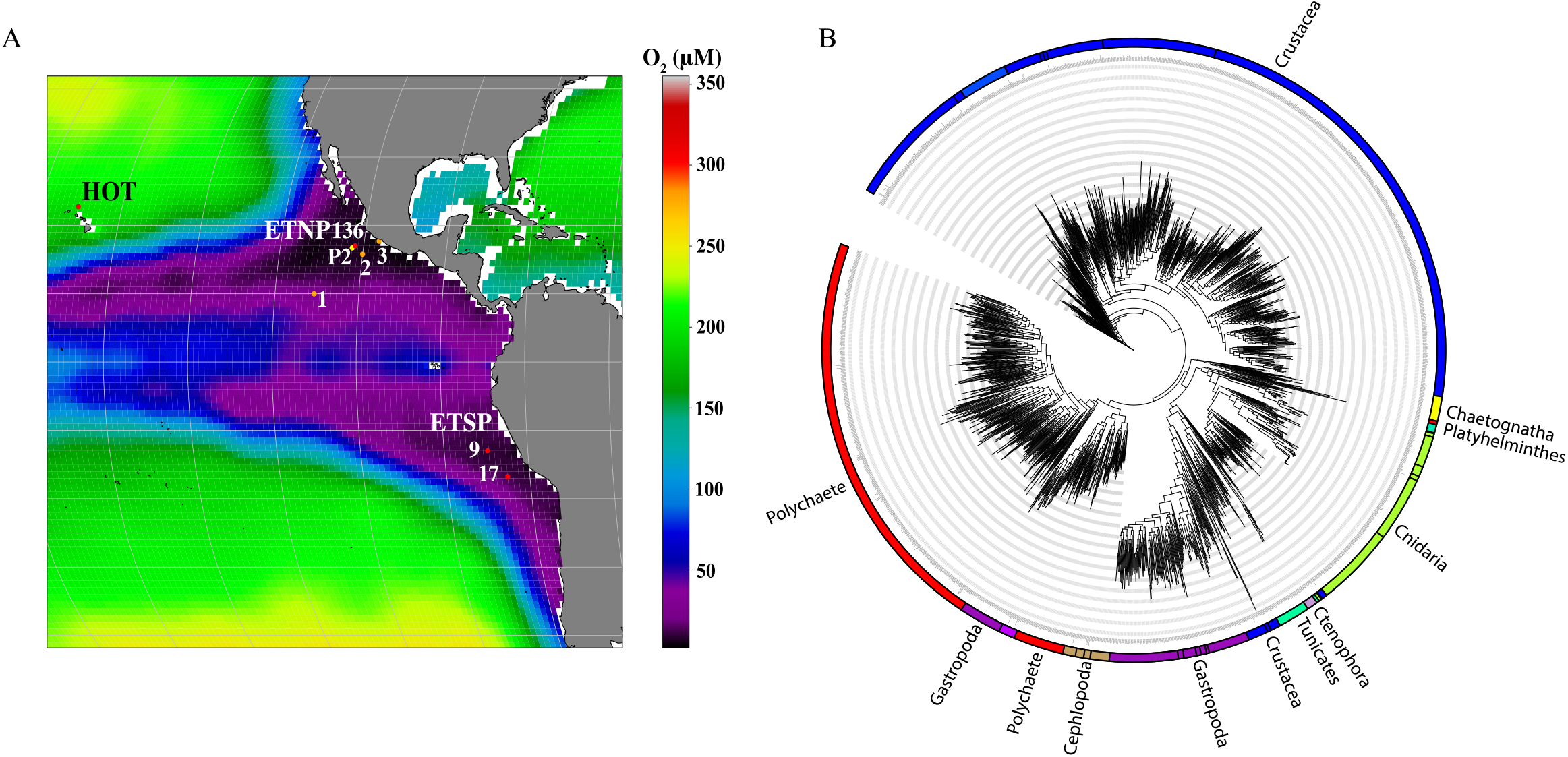
A) Map of stations with metagenomic depth profiles (red circles), eukaryotic metatranscriptomes (orange circles) and St P2 (yellow circle). Background colors indicate oxygen concentrations at 250m as found in World Ocean Atlas 2018 (Boyer et al 2018). B) Cytochrome c oxidase gene (COI) nucleotide phylogenetic tree for zooplankton. A more detailed version of this tree can be seen in supplemental figure 1.

### Metagenomes

Cellular metagenomes (>0.2 µm) from the Eastern Tropical South Pacific St 9 (13ºS and 82.2ºW) and St 17 (16.7ºS and 79ºW) can be downloaded from Bioproject PRJNA704804 [53]. Hydrographic and nutrient data from this July 2013 ETSP cruise are deposited at NODC as accession 0128141 and are previously published [53–55]. Cellular metagenomes from the Eastern Tropical North Pacific in 2012 St 136 (17.04 ºN, 106.54 ºW) can be downloaded from Bioproject PRJNA350692 [7]. Hydrographic and nutrient data from this April 2012 ETNP cruise are deposited at the NODC as accession 0109846. Data for ETNP St 136 can be seen in [7]. Hawaii Ocean Time-series (HOT) (22º45’N and 158ºW) cellular metagenomes from May (HOT 272), August (HOT 275), and November (HOT 278) of 2015 [56, 57] were downloaded from Bioproject PRJNA352737. Nutrient and CTD measurements for these cruises can be found at Hawaii Ocean Time Series Data Organization and Graphical System (https://hahana.soest.hawaii.edu/hot/hot-dogs/).

ESTP assembled contigs have not been previously published and have been uploaded to NCBI WGS under Bioproject PRJNA704804. Contigs were assembled with MEGAHIT (v 1.2.8) [58] and annotated with PROKKA [59]. Translated proteins were made into a custom blast database, which was then queried for urease (*ureC*), and taurine metabolism proteins. These assembled proteins were used in many of the phylogenetic trees described below.

### Phylogenetic trees and metagenomic read placement

COI gene sequences (∼650 bp) were downloaded from the Census of Marine Zooplankton [52]. After dereplication, these 650 bp sequences were aligned using MUSCLE [60] to construct a nucleotide maximum likelihood phylogenetic tree using RaxML-ng with bootstrap analysis (n=100) [61]. Groups within the phylogenetic trees were then labeled based on the references within that group (Figures 1B and S1).

The sequences making up the tree were then compared against ETNP, ETSP, and HOT metagenomic read databases using tblastn with an e-value of 10e-5. The short reads were then aligned to the reference tree using PaPaRa Parsimony-based Phylogeny-Aware Read Alignment program 2.0 [62]. Non-overlapping paired-end reads were then combined into one aligned sequence and placed on the tree by EPA-ng [63]. Placed reads have a pendant length indicating the similarity between a query read and the location it places on the tree. Reads that were placed with a pendant length greater than 2 were removed. The remaining reads were enumerated for each taxonomic group using the “assign” subcommand of Gappa [64]. Taxonomic read counts were normalized using the method previously described by [20] where normalization factors for each sample were determined by dividing the number of good quality reads in a sample by the 100 m ETNP sample. The read counts were multiplied by the sample normalization factor, divided by the gene length, and then multiplied by 100 to make visualization easier. A comparison of raw read counts and normalized read counts for zooplankton groups can be seen in Table S2.

A similar analysis with a phylogenetic tree made of Ichthyoplankton (fish) COI genes from the Census of Marine Zooplankton [52] was unsuccessful due to the small numbers of Ichthyoplankton reads in the metagenomes (data not shown).

The microbial community was analyzed using phylogenetic read placement of the RNA polymerase gene (*rpoB*) as above. The protein phylogenetic tree for RNA polymerase gene (*rpoB*) from [7] and the protein urease (*ureC*) phylogenetic tree from [51], which both included reference sequences and ETNP contigs, were updated using single cells [65] and environmental contigs from marine oxic environments [66] and the ETSP (Fig S2-3). Taurine metabolism genes [50], a substrate-binding subunit of ABC transporter *tauA*, organic S assimilation taurine dioxygenase gene *tauD*, and energy providing C and N assimilation taurine:pyruvate transaminase gene *tpa*, were created using the same set of contigs (Fig S4). Proteins from [49] were used to validate the assignment of proteins on taurine metabolism trees. For example, other proteins were often misannotated as *tauD*, and *tpa* proteins were often misannotated as other proteins. The % of the prokaryotic community was calculated by dividing the normalized reads of a particular organism by the total bacteria and archaea *rpoB* reads. The % of SAR11 was calculated by dividing the normalized reads of a gene assigned to SAR11 by the *rpoB* reads for SAR11.

The abundance of eukaryotic algae numbers were taken directly from [53], where they were obtained by using phylogenetic read placement of the 18S rRNA gene on a tree constructed from sequences encompassing the PR2 database [67].

### Suspended Particulate Organic Carbon and Nitrogen and other chemical analyses

In ETSP 2013 and ETNP 2018, water for suspended particulate organic C and N analyses was obtained from Niskin bottles and filtered onto pre-combusted GF/F filters. At 6 stations in the ETSP 2013, either 10 L or 20 L were filtered and then split into four. One-fourth was used for POM analysis. The latitude and longitude for all 6 ETSP stations can be found in Fig S5. Each station had samples from 2-6 depths. Therefore, we combined stations from the same ODZ regimes for graphing purposes. In the ETNP St P2 (16.9ºN 107ºW) (Figure 1) from May 2018, either 4L or 10L (in deep samples) were filtered at 18 depths and the entire sample was utilized for POM. In ETNP St P2 2012, water was instead filtered onto 142 mm pre-combusted GF/F filters *in situ* using a series of McClane pumps (on average ∼300-500 L). Two punches from a size 12 core bore punch (21 mm diameter) were obtained from each filter and used for POM analysis. In all cases, samples were wafted with HCl overnight to remove carbonate, dried at 40°C, and sent to UC Davis Stable Isotope Facility for C and N analysis. All POM data can be seen in Table S3.

July 2013 monthly MODIS R2018 satellite chlorophyll data was downloaded from the Oregon State Ocean Productivity webpage. Surface chlorophyll for ETSP stations was extracted from a 1/3° box around each station.

Biological N_2_ gas concentrations were sampled at P2 in April 2012 and were analyzed exactly as in [29]. To obtain biological N_2_ gas concentrations, background N_2_ gas from air equilibration and mixing was removed [29].

An ADCP-75 was used continuously during the April 2012 ETNP cruise to track micronekton migration. Data was analyzed with a publically available R script ADCP_PlotTimeseries_R.pynb, and St P2 is shown here.

### qPCR of metazoan DNA

Duplicate eDNA samples were obtained at ETNP St P2 (16.9ºN 107ºW) (Figure 1) in October 2019 on the RV *Kilo Moana* by vacuum filtering ∼4 L of water onto 47 mm diameter, 1 µm pore size cellulose nitrate membrane filters (Whatman). Water for duplicates was obtained from separate Niskin bottles on the same rosette. Samples were obtained during midday and ∼midnight on two separate days. DNA was extracted using a CTAB Chloroform/Isoamyl extraction method [68]. To create a metazoan DNA standard, a zooplankton net tow was performed with a 64 µm mesh at the Horn Point dock in the Choptank River, Chesapeake Bay MD, USA. DNA was immediately extracted from a portion of the zooplankton tissue mass via the EZNA tissue kit (Omega Biotek) and was used to create a highly concentrated standard whose DNA concentration was measured with a Qubit 2.0 (ThermoFisher). qPCR for the COI gene was performed using Leray primers (mlCOIintF: GGWACWGGWTGAACWGTWTAYCCYCC and jgHCO2198: TAIACYTCIGGRTGICCRAARAAYCA) [69] and SsoAdvanced Universal SYBR Green Supermix (Bio-Rad) on a Bio-Rad C1000 Touch Thermal Cycler with a Bio-Rad CFX96 Touch Real-Time PCR Detection System for 45 cycles. Each sample was run in triplicate. Two standard curves (10,500 pg/uL, 1,050 pg/uL, 105 pg/uL, 10.5 pg/uL, and 1.05 pg/uL) were run along with the samples. Metazoan eDNA concentrations from the depth profile were reported in ng/L based on the standard curve of coastal zooplankton DNA.

### Eukaryotic transcriptomes

Onboard R/V *Sally Ride* in March/April 2018, three stations in the eastern tropical North Pacific were visited for transcript sample collection (Figure 1). Station 1 (10°N 113°W) was an offshore station; Station 2 (15.77°N 105°W) was located in the heart of ODZ; Station 3 (17.68°N 102.35°W) was a coastal station. Dissolved oxygen concentration was determined using the SBE 43 dissolved oxygen sensor attached to an SBE 911+ Conductivity, Temperature, and Depth (CTD) system. A diaphragm pump (Husky Model 307) connected to a 142-mm stainless steel filter holder was used to filter 28.5 L of seawater onto a 0.2 μm polycarbonate filter. No organisms were visible on the filters. Each filter was immediately folded, placed in a 2-ml externally threaded cryogenic vial, and flash-frozen in liquid nitrogen before storage at −80°C. Samples were shipped to the laboratory on land on dry ice.

In the laboratory, total RNA was extracted using the AllPrep DNA/RNA/miRNA Universal Kit (QIAGEN) following the AllPrep DNA/RNA/miRNA Universal Handbook except that the cell disruption step was customized for our samples. Filters were first cut into 2 mm by 2 mm pieces using scissors sterilized and treated with RNase AWAY surface decontaminant (Thermo Scientific). Filter pieces were transferred into MN Bead Tubes Type A (Macherey-Nagel) containing 800 μl of buffer RLT Plus that were pre-chilled at −80°C for 10 minutes. Bead beating of the samples was performed for 60 seconds using a Bio-spec Mini-BeadBeater-16. The remaining nucleic acid extraction steps followed the AllPrep DNA/RNA/miRNA Universal Handbook without modifications. An extraction blank was included for each batch of extraction procedures. RNA yield was quantified using a Qubit fluorometer (ThermoFisher Scientific) following the manufacturer’s instructions. RNA quality was determined using a TapeStation 2200 system (Agilent) and a NanoDrop 1000 spectrophotometer (Thermo Fisher Scientific). All RNA samples were shipped on dry ice to the DNA Technologies Core at the University of California Davis for library construction with poly-A enrichment and sequencing on one lane of the NovaSeq 6000 platform (Illumina) with S4 type flow cell (150 bp PE).

The average number of raw reads for the 32 polyA-enriched metatranscriptome libraries was 13,695,246. The number of raw reads for each metatranscriptome can be seen in Supplementary Table S4. Raw reads were quality filtered using the BBDuk tool in the BBMap software package (v38.87) with the parameters “minlen=51 minlenfraction=0.33” and Trimmomatic [70] (v0.39) with the parameters “ILLUMINACLIP:2:30:10 LEADING:3 TRAILING:3 SLIDINGWINDOW:4:15 MINLEN:100”. Ribosomal RNA and internal transcribed spacer (ITS) regions were removed with SortMeRNA [71] (v4.2.0) using the SILVA database [72] (Release 138.1) for large and small subunit rRNA, the Rfam database 14 [73] for 5S and 5.8S rRNA, and the UNITE database (Release 2021.5.10) [74] for eukaryotic ITS as references. Filtered RNA reads from 32 samples were co-assembled using MEGAHIT [58] (v1.2.9), resulting in 7,649,133 contigs with a total length of 3,971,957,536 bp. Protein-coding regions of the co-assembled metatranscriptome were identified *ab initio* by GeneMarkS-T [75] and annotated by InterProScan [76] (v5.51-85.0). The metatranscriptomic contigs were queried against the NCBI nr database using DIAMOND [77] with an e-value threshold of 1×10^−5^ and the options “--sensitive--min-orf 20”. The resultant NCBI taxonomy ID was used to assign taxonomy to each contig. The filtered reads were mapped to the metatranscriptomic contigs using bowtie2 (v.2.4.2) [78] and the number of reads mapped to protein-coding regions were determined with featureCounts (v2.0.2) [79] using the parameters “--fracOverlap 0.5 -M -- fraction --donotsort”. Normalized reads per kilobase of transcript per million reads mapped (RPKM) was computed with EdgeR [80], which accounted for both the size of the metatranscriptome library and the length of the protein-coding region.

### Cnidaria photos

Underwater Vision Profiler-5 (UVP) data from ETNP St P2 in January 2017 on the RV *Sikuliaq* was published in [81]. The UVP, which optically examines 1 L at a time, was attached to the CTD rosette during all casts. UVP photos from ETNP St P2 were examined manually for Cnidaria.

## Results

### Organic Matter

We examined whether zooplankton could be affecting the organic matter composition in the ODZ. We compare profiles at P2 between years, but the ADCP backscattering profiles stayed consistent at this station over several cruises in the [21] dataset and over the four cruises represented here (2012, 2017, 2018, 2019) (data not shown). The EK60 backscattering (3800 Hz) maximum vertical migration depth for ETNP station P2 was 270m-400m, with a distinct peak at 260-280m (Fig 2C; [81]). A similar maximum in ADCP backscattering was found at the same depths at St P2 in 2012 (Fig S6). ETNP St P2 from 2018 had a detailed POM profile across this 260-280m backscattering maximum (Fig 2B, 2C). C:N ratios are a way to examine whether the organic matter is fresh or degraded, with high C:N ratios (>9) indicating degraded material.

**Figure 2.**
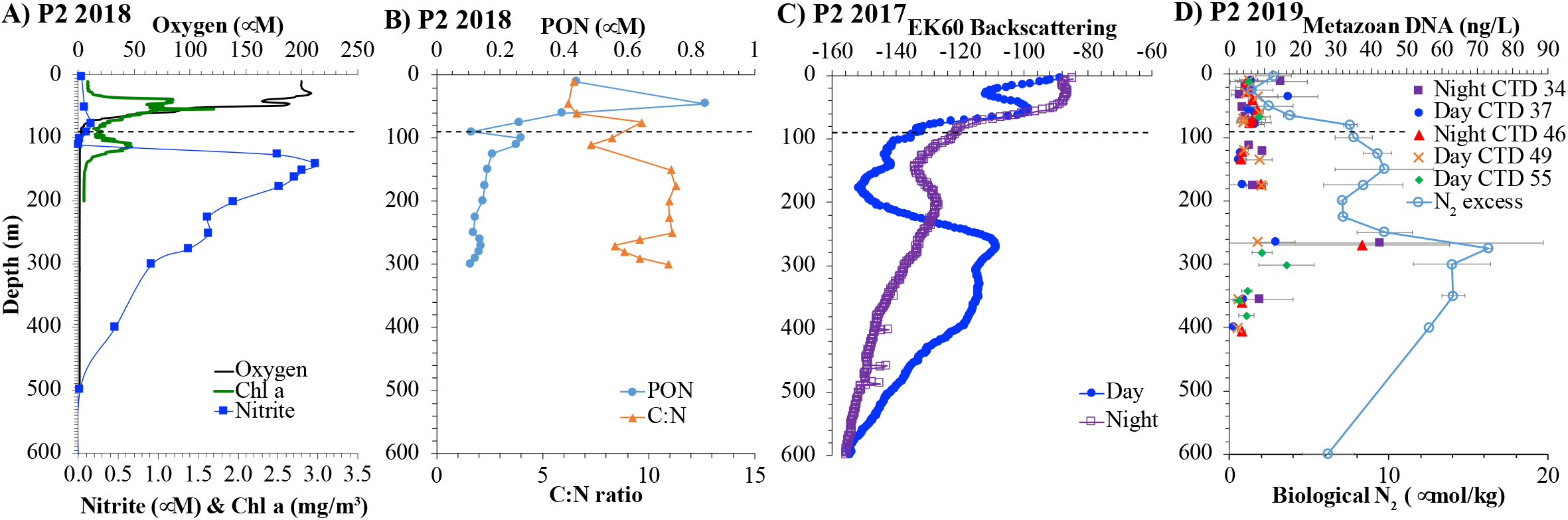
A) Oxygen, chlorophyll fluorescence, and nitrite concentrations from ETNP St P2 2018, B) Suspended particulate organic nitrogen and C:N ratios of suspended particulate organic matter from ETNP St P2 2018, C) EK60 Backscattering data (3800 Hz) from ETNP St P2 2017 reproduced from (Cram et al 2022), and D) Metazoan DNA concentrations from qPCR from ETNP St P2 2019, where the standard was quantified DNA from a coastal net tow. Error bars in DNA concentrations represent differences between water from different Niskin bottles closed at the same depth. Also, biological N_2_ gas from St P2 in 2012 is shown.

Additionally, different sources of organic matter have different C:N signatures. Phytoplankton C:N values of ∼6 are higher than crustacean zooplankton of C:N values of 4.8-6.2 [82]. Jellyfish are also protein-rich with a C:N ratio of 4.5:1 [39]. In the euphotic zone at St P2, the C:N ratio of suspended organic matter was 6, consistent with the presence of phytoplankton, but increased to 10 in the oxycline, where concentrations of C and N had a minimum; then C:N decreased to 7 at the secondary chlorophyll maximum (Figure 2AB, S7), which is dominated by *Prochlorococcus* cyanobacteria [20]. Starting at 150m, the C:N ratio was 11 for most of the deep water, indicating degraded material, but had a smooth decrease to 8 at 270m, then increased again to 11 (Figure 2B). qPCR data of metazoan DNA had a huge maximum at 270m at this station (Figure 2D). The highest yields of DNA per liter of seawater were from night casts, however, the error bars, representing water from different Niskin bottles at the same depth, are too large to definitively indicate diel cycling (Figure 2D). The high quantity of metazoan DNA at 270m is directly coincident with the low C:N ratios of suspended POM at 270m. Additionally, both are coincident with a small nitrite maximum (Figure 2A) and the highest biological N_2_ gas concentration (Figure 2D). Similarly, in suspended particulate organic matter from St P2 in 2012, there is a shift in C:N ratios from ∼8.5 in most of the ODZ to 6 at a single point at 250m (Fig S6). While generally less degraded in April 2012 than May 2018, the C:N ratios follow a similar pattern with lower C:N ratios at the vertical migration (backscattering) maximum.

Depleted δ^13^C values (−27‰ to −30‰) are indicative of slow-growing small-celled phytoplankton while fast-growing large-celled phytoplankton have δ^13^C values ∼-20‰ [83]. The δ^13^C varied between −24‰ and −27‰ at St P2 in 2018 with the most depleted values at the bottom of the primary chlorophyll maximum and the secondary chlorophyll maximum (Figure S6). δ^13^C of suspended particulate organic C were more enriched at the vertical migration depths (−24‰; Figure S7). However, this enrichment is within the normal bounds of algae δ^13^C values.

Suspended POM concentrations were also obtained in the ETSP in 2013, though only a few depths were filtered at each station. ETSP St 9 and St 17 had >4000m water depth and were neither coastal nor far offshore. Satellite chlorophyll data indicated similar amounts of surface chlorophyll at ETSP St 9 and St 17 during the cruise in July 2013, but twice as much chlorophyll at coastal station BB2 (1600m bottom water depth) and more reduced chlorophyll at more offshore stations (Fig S5). Satellite chlorophyll from the previous month (June), indicates that St 17 had higher surface chlorophyll than St 9 due to a bloom reaching out from the coast (Fig S5). However, CTD data at the exact time of sampling indicates that the chlorophyll maximum at St 9 was twice as large as seen at St 17 and the nitrite maximum was also much larger at St 9 (Fig 3, S8). Thus, the amount of surface productivity at these stations is likely variable. At the time of sampling, the amount of suspended organic C at depth at Stations 9 and 17 were similar to each other and to concentrations at the BB2 coastal station (Figure S9). However, offshore stations had significantly less organic C at depth (Figure S9). Contrastingly, the C:N ratios at St 9 and St 17 were quite different. St 9 and coastal station BB2 both had secondary chlorophyll maxima and C:N ratios ∼6-8 at depth (Figure S9). However, St 17 did not have a secondary chlorophyll maximum and had high C:N ratios of ∼12-15 at depth, indicating degraded organic matter, similar to the offshore stations (Figure S9). Thus, more organic N was available to microbes at St 9 than at St 17. Additionally, the δ^13^C of suspended POC at St 9 and coastal station BB2 were − 22‰ to −24‰, usual values for coastal phytoplankton communities, while the δ^13^C of St 17 and the offshore stations were −24‰ to −26‰ (Figure S9), indicative of slow-growing small celled phytoplankton usually found offshore [83]. Thus, at the time of sampling, St 17 was more similar to offshore stations than was St 9 but still retained a significant ODZ and nitrite maximum.

**Figure 3.**
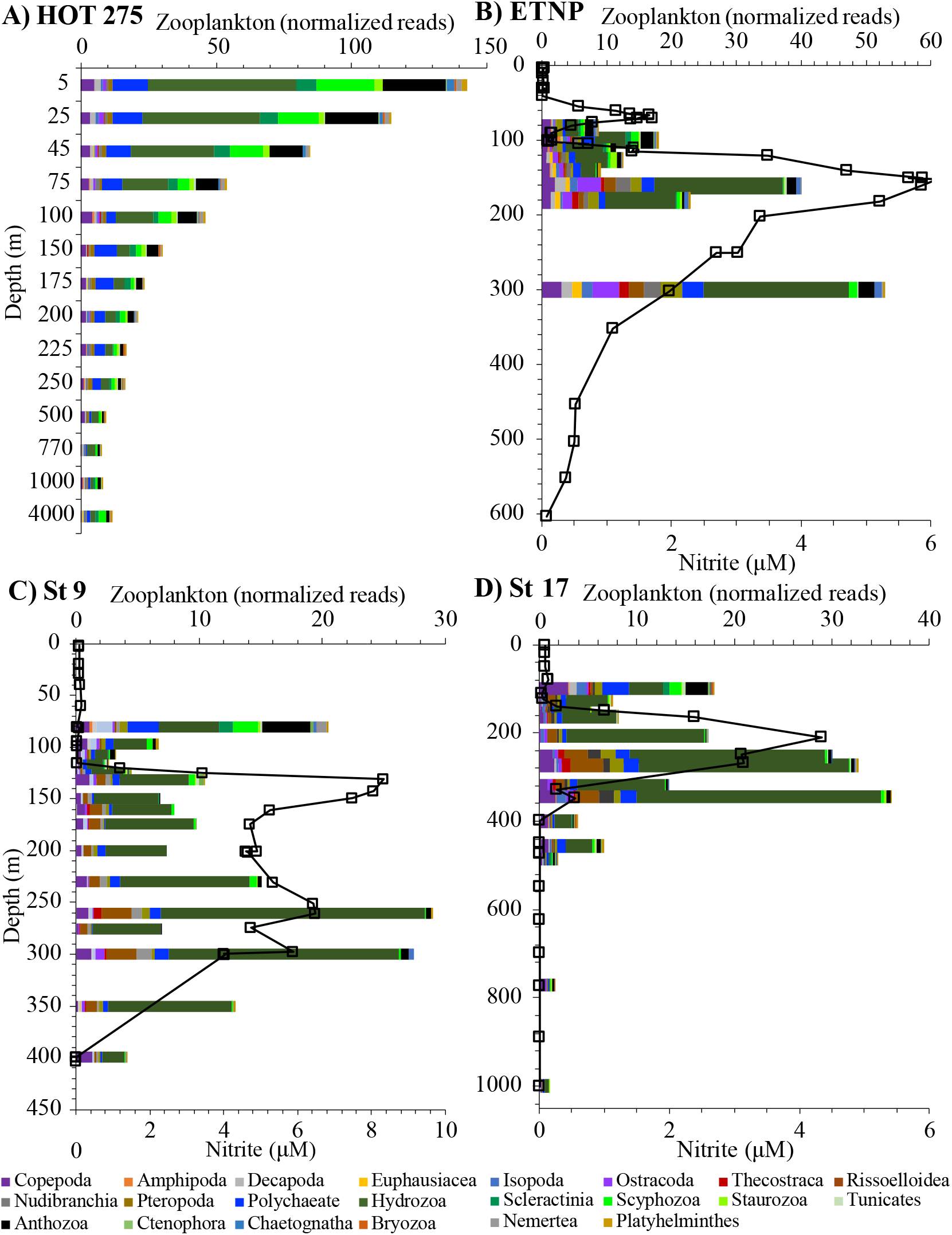
Zooplankton groups from metagenomic data for the COI gene examined by normalized reads for A) HOT 275, B) ETNP, C) ETSP St 9, and D) ETSP St 17. Squares indicate nitrite concentrations at the same stations. Data as % of zooplankton is in Fig S11.

Suspended Particulate C concentrations (0.7 µm pore size filter) from HOT in 2015 include both organic and inorganic C [56, 84]. Maximal values were between 2-3 µM C (Figure S10). In HOT 272, the maximum extended from the surface to 175m (Figure S10A). In HOT 275, the maximum was at 50m (Figure S10B). In HOT 278, the maximum was in the mixed layer (0-56m) (Figure S10C).

### Metagenomics

HOT metagenomes had a relatively smooth decrease in zooplankton normalized reads with depth (Fig 3, S11). Cnidaria dominated HOT profiles (40-75% of zooplankton reads) with Hydrozoa (averaging ∼25% of zooplankton reads), and Scyphozoa (true jellyfish; ∼10% of zooplankton reads), as the most abundant groups (Fig 3, S11). Polychaetes were also fairly abundant (averaging 11-15% of zooplankton reads) at HOT (Fig 3, S11). Crustaceans (13-15% of zooplankton reads) were dominated by copepods (∼50% of crustacean reads) and decapods (∼20% of crustacean reads) (Fig 3, S11). HOT 272 770m had an overabundance of copepods (>50% of zooplankton reads), but this was not seen in other months making us wonder if a carcass was caught on the filter (Fig S11).

At the ODZ stations, which included zooplankton vertical migration depths, the zooplankton normalized reads were much more variable between depths (Fig 3). The fully oxic depths in the ETNP resembled HOT profiles. However, Polychaetes were lower in the ODZs (5-7% of zooplankton reads) (Fig 3, S12). On the other hand, Gastropods (Rissoelloidea, Nudibranchia, and pteropods) were more prevalent in all three ODZ stations (averaging ∼11% of zooplankton reads) than at HOT (∼6% of reads) (Fig 3, S12). Crustaceans such as ostracods and euphausiids had increased prevalence in the ETNP ODZ at both 160m and 300m as did pteropods and Rissoelloidea (Fig 3, S12). In the ETNP, the 300m sample was the only metagenomic sample in the backscattering zone of strong vertical migration. The 300m metagenomic sample did have the most zooplankton DNA of the depths sampled, but the 160m sample also had a lot of zooplankton DNA (Fig 3), indicating that the backscattering maximum did not account for all zooplankton. Backscattering measurements preferentially see large zooplankton or fish and animals with swim bladders, air pockets, or shells [85]. Thus, not all vertical migrators are seen by backscattering.

Hydrozoa averaged 35-60% of zooplankton reads and had multiple maxima in the ODZs (Fig 3, S12). To get more evidence for Hydrozoa in the ODZ, we manually examined UVP photographs from ETNP St P2. In this qualitative analysis, we found several <5 mm jellyfish both in and below the ODZ (Figure S13). Since it only examines 1 L at a time, the UVP is biased toward small organisms. However, it shows that these tiny jellyfish do exist in the ODZ.

Multiple nitrite maxima were found in the ETSP ODZs (Fig 3). The smaller deeper nitrite maxima may be coincident with higher numbers of zooplankton normalized reads (Fig 3).

### Eukaryotic transcripts

Because no animals were visible on the transcript filters, we believe that these eukaryotic transcripts (polyA tail selected) of metazoans are eRNA, similar to eDNA. As transcripts degrade quickly, this eRNA indicates sources of fresh organic matter. The most active metazoan taxa in the ETNP included Tunicata, Copepoda, Hydrozoa, Anthozoa, Scleractinia, and Platyhelminthes (Figure 4). The activity of Hydrozoa, Anthozoa, Scleractinia, (all Cnidaria) and Platyhelminthes were higher in the oxycline than in the fully oxygenated surface water. However, the general activity was much reduced in the ODZ, and tunicates were disproportionately missing (Figure 4). The overall activity of zooplankton groups in the ETNP was lower at Station 1 (offshore station) than at Station 2 (ODZ core station; student’s t-test p-value = 0.018) and at Station 3 (coastal station; student’s t-test p-value = 0.035). Stations 2 and 3 had secondary nitrite maxima indicating anoxia, but Station 1 did not have a secondary nitrite maximum and was likely suboxic rather than truly anoxic Fig S14 [86].

**Figure 4.**
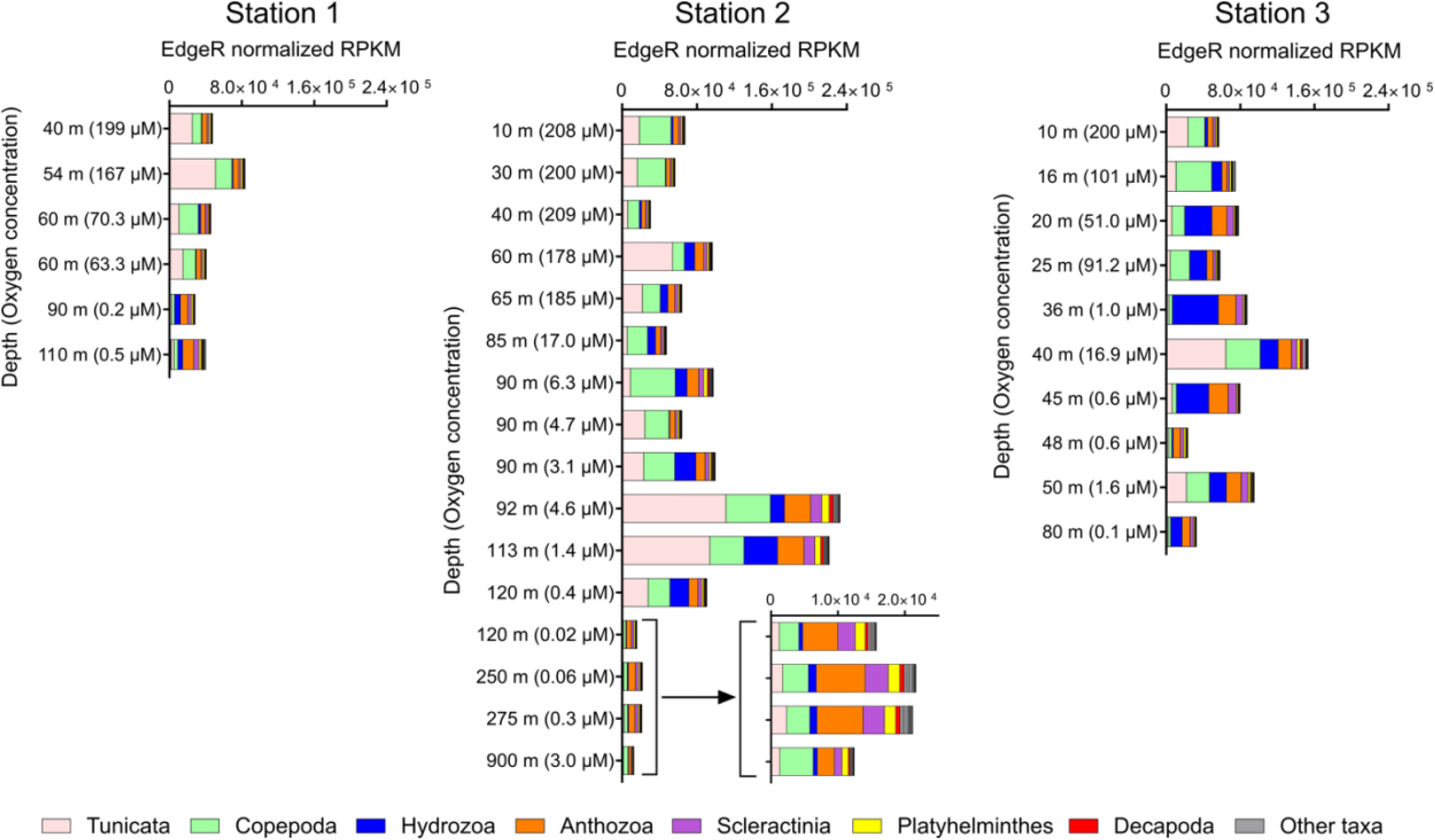
Metazoan eRNA transcripts for ETNP stations 1, 2, and 3 from 2018. Depths at each station are listed with oxygen concentrations in parentheses. Oxygen below 1 µM can be considered below detection for a Seabird 43 dissolved oxygen sensor.

### SAR11

The ETNP (28-60%) and HOT (20-35% of the prokaryotic community) had higher proportions of SAR11 in the microbial community compared to the two ETSP stations (8-27%) (Fig S15). At all four stations, SAR11 normalized reads correlated strongly (generally R^2^ >0.8) with total zooplankton COI normalized reads, and these correlations were particularly high in the ODZs (Table 1). At HOT, both SAR11 and zooplankton had smooth depth profiles decreasing with depth (Figure 5 and S16). In the ODZs, SAR11 bacteria do not have smooth depth profiles, but rather have many maxima (Figure 5). These maxima match maxima in Cnidaria, gastropod, and crustacean depth profiles (Figure 5). The R^2^ between SAR11 and Cnidaria profiles were high at all stations and reached 0.98 (p=0.0002) in the ETNP ODZ (Table 1). The R^2^ between SAR11 and Gastropoda was high in the ODZs, reaching >0.8 in both the ETSP St 17 and ETNP, but was generally not significant at HOT (Table 1). Crustaceans and SAR11 were not significant at ETSP St 9 (Table 1). Correlations between SAR11 and cyanobacteria were not significant, and correlations between SAR11 bacteria and eukaryotic algae were much weaker and inconsistent between seasons (Table S5; Fig S15 and S17). However, HOT 272, the May sampling period, did have significant correlations with eukaryotic phytoplankton (R^2^=0.55; p=0.006) (Table S5; Figure S17). Similarly, HOT 272 was the only season to correlate with suspended particulate carbon and nitrogen concentrations (R^2^= 0.61 and 0.53; p= 0.01 and 0.03) (Table S5).

**Table 1.** Correlations between SAR11 (*rpoB*) normalized reads and Zooplankton (COI) normalized reads. Inter = Intercept. Pval=P value.

|  | HOT 272 | HOT 275 | HOT 278 | ETNP | ETNP ODZ | ETSP 17 | ETSP 17 ODZ | ETSP 9 | ETSP 9 ODZ |
| --- | --- | --- | --- | --- | --- | --- | --- | --- | --- |
| **Total Zooplankton** |  |  |  |  |  |  |  |  |  |
| slope | 0.63±0.07 | 0.42±0.06 | 0.4±0.1 | 0.05±0.01 | 0.077±0.007 | 0.22±0.02 | 0.23±0.05 | 0.11±0.01 | 0.11±0.01 |
| Inter. | -117±20 | -67±17 | -72±30 | 6±5 | -4±3 | -3±2 | -3±4 | 2±1.7 | 2±1.8 |
| R<sup>2</sup> | 0.89 | 0.85 | 0.59 | 0.63 | 0.96 | 0.86 | 0.80 | 0.73 | 0.74 |
| P val | 0.000003 | 0.00005 | 0.003 | 0.01 | 0.0005 | 0.000002 | 0.003 | 0.00002 | 0.00009 |
| **Cnidaria** |  |  |  |  |  |  |  |  |  |
| slope | 0.47±0.05 | 0.33±0.04 | 0.32±0.08 | 0.08±0.01 | 0.102±0.007 | 0.4±0.06 | 0.41±0.08 | 0.19±0.03 | 0.19±0.03 |
| Inter. | -90±15 | -59±12 | -54±22 | 0±4 | -7±3 | -8±5 | -4±9 | 3±3 | 3±3 |
| R<sup>2</sup> | 0.89 | 0.87 | 0.58 | 0.88 | 0.98 | 0.77 | 0.78 | 0.68 | 0.69 |
| P val | 0.000004 | 0.00002 | 0.004 | 0.0001 | 0.00002 | 0.00003 | 0.003 | 0.0001 | 0.0002 |
| **Crustacea** |  |  |  |  |  |  |  |  |  |
| slope | 0.039±0.008 | 0.022±0.006 | 0.038±0.008 | 0.016±0.003 | 0.019±0.002 | 0.021±0.006 | 0.021±0.006 |  |  |
| Inter. | -5±2 | -1±2 | -4±2 | 0.6±0.9 | -0.8±1 | 0.1±0.5 | -0.2±0.5 |  |  |
| R<sup>2</sup> | 0.68 | 0.60 | 0.68 | 0.82 | 0.90 | 0.50 | 0.70 | 0.21 | 0.28 |
| P val | 0.001 | 0.005 | 0.001 | 0.0008 | 0.001 |  |  | 0.07 | 0.052 |
| **Gastropoda** |  |  |  |  |  |  |  |  |  |
| slope |  |  | 0.014±0.004 | 0.010±0.001 | 0.012±0.001 | 0.036±0.005 | 0.040±0.007 | 0.014±0.002 | 0.014±0.002 |
| Inter. |  |  | -1±1 | 0.1±0.5 | -0.8±0.5 | -0.88±0.4 | -1.1±0.8 | 0.2±0.2 | 0.2±0.2 |
| R<sup>2</sup> | 0.05 | 0.28 | 0.49 | 0.87 | 0.94 | 0.81 | 0.82 | 0.67 | 0.69 |
| P val | 0.4 | 0.09 | 0.01 | 0.0002 | 0.0003 | 0.00001 | 0.001 | 0.00009 | 0.0002 |

**Figure 5.**
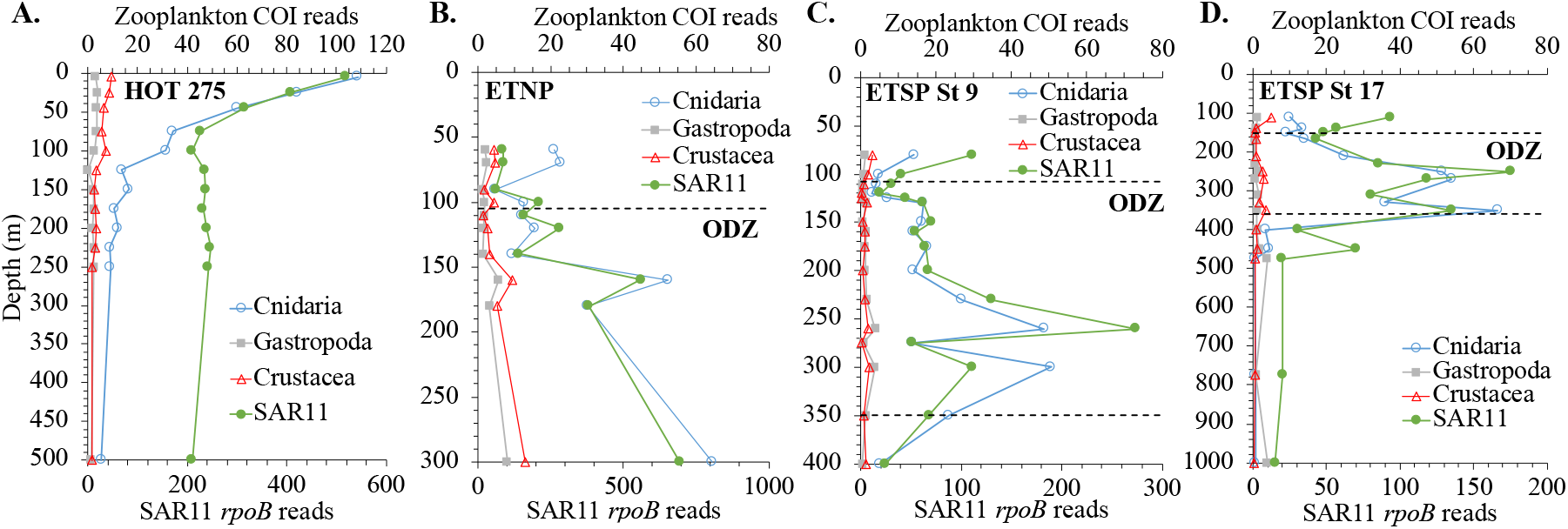
Depth profiles of Cnidaria, Gastropoda, and Crustacea (COI) and SAR11 (*rpoB*) metagenomic reads in the A) HOT 275, B) ETNP St 136, C) ETSP St 9, and D) ETSP St 17.

One way that zooplankton could feed free-living SAR11 bacteria would be with their excreta. Crustacean zooplankton and at least some Gastropods excrete urea [37]. At HOT in August (275) and November (278), ∼20% of SAR11 bacteria contained a urease gene in the upper euphotic zone, and in May (272), 30% of SAR11 bacteria contained a urease gene (Fig 6). The SAR11 urease gene was found at two different sections of the phylogenetic tree separating into oxic and ODZ phylotypes (Fig S3) [51]. In the ETNP ODZ, only 10% of SAR11 had the urease gene and in the ETSP stations, only 5% of SAR11 had the urease gene (Fig 6). However, this is in contrast to similar depths at HOT where no SAR11 was found (Fig 6).

**Figure 6.**
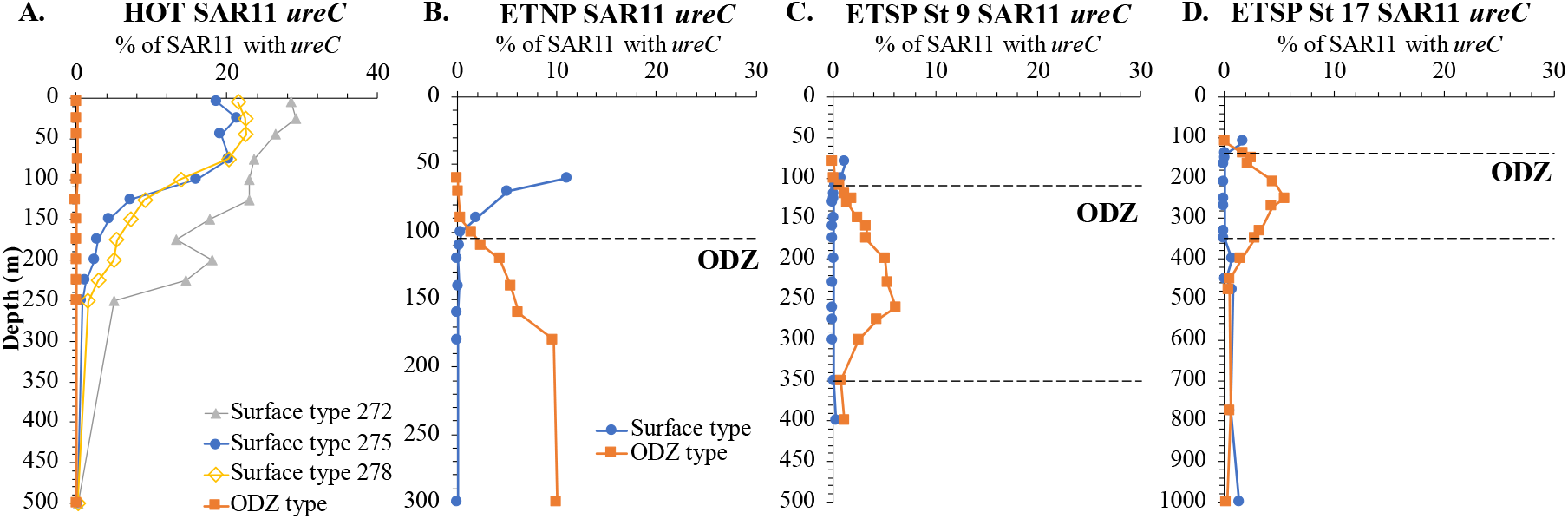
SAR11 urease (*ureC*) metagenomic depth profiles shown in % of SAR11 with *ureC* at

Both crustaceans and Cnidaria excrete taurine [35]. Taurine metabolism in SAR11 includes two separate pathways [49, 50]. The tau ABC transporter (including substrate-binding subunit *tauA*) provides taurine to both an organic S assimilation pathway (here represented by taurine dioxygenase gene *tauD*), and a C and N assimilation pathway (here represented by taurine: pyruvate transaminase gene *tpa*). The gene *tpa* seems to be ∼1 copy per SAR11 genome throughout our dataset, perhaps increasing to 1.5 copies in the ETSP ODZs (Fig 7, S18). However, the % of SAR11 with *tauD* was reduced both in surface waters and in the ODZs but was elevated in deep oxic and hypoxic waters (Fig 7, S18). Transporter *tauA* was generally ∼1 copy per SAR11 genome but became elevated in the hypoxic waters underneath the ODZ at ETSP St 17 (Fig 7). Thus, all the SAR11 could use taurine, but many SAR11 in the ODZ had lost the ability to use taurine for S assimilation but retained the ability to use taurine for C and N assimilation. On the other hand, taurine may be an important source of organic S in the deep hypoxic and oxic ocean as some SAR11 appeared to have more than 1 copy of *tauD* in deep waters.

**Figure 7.**
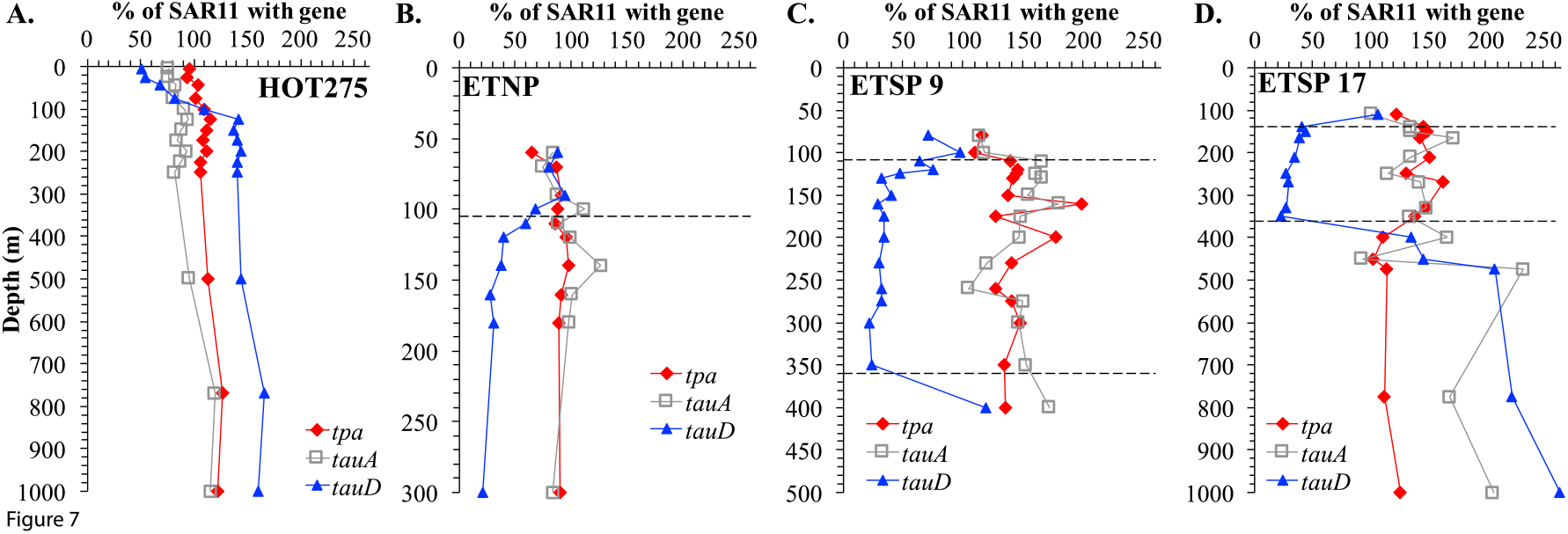
Metagenomic depth profiles of taurine metabolism genes taurine:pyruvate transaminase (*tpa*), taurine dioxygenase (*tauD*), taurine ABC transporter substrate binding subunit (*tauA*) in SAR11 for A) HOT 275 (Aug), B) ETNP, C) ETSP St 9. D) ETSP St 17. The dashed lines indicate the boundaries of the ODZ. A percentage >100% indicates multiple copies of the gene.

## Discussion

We find clear evidence that zooplankton affect the organic matter at a vertical migration depth in the ODZ. The EK60 backscattering maxima at ETNP St P2 encompasses 260-400m with a distinct peak at 260-280m (Fig 2). The shape of this peak was reproducible between ADCP and EK60 backscattering data and between years at the same station (Fig 2, S6). The well-defined suspended organic matter C:N minimum at 270m was reproducible between 2012 and 2018 (Fig 2, S6) and was coincident with a qPCR maximum in metazoan DNA (Fig 2). Crustacean zooplankton have C/N values of 4.8-6.2 [82] and jellyfish have a C:N ratio of 4.5:1 [39], both of which are lower than phytoplankton values ∼6. Thus, this C:N ratio minimum is likely due to migrating zooplankton. No zooplankton bodies were seen on the POM filters, rather zooplankton can create suspended POM by molting or defecating [87]. This data is consistent with previous UVP particle measurements at ETNP St P2 that suggest small increases in particle flux in the region of diel migration in the ODZ [81]. Additionally, a small nitrite sub-maximum and the highest biological N_2_ gas concentration are coincident with maxima in these other parameters between 260-280m (Fig 2). The introduction of labile organic matter by zooplankton could be stimulating nitrate reduction and N_2_ production at that depth. Similarly, at ETSP St 9, nitrite concentrations have three maxima and the second and third peaks are coincident with maxima in zooplankton DNA in metagenomes (Fig 3), and ETSP St 17 has two nitrite maxima and the second peak corresponds to a maximum in zooplankton DNA in metagenomes (Fig 3). Together these data indicate the transfer of organic matter from zooplankton to bacteria in the water column.

Examining zooplankton through metagenomics has two advantages over eDNA amplicons. First, PCR-based analyses are biased, both over and under amplifying some organisms [88]. The COI gene is not conserved enough for one set of primers to amplify all zooplankton [89]. Second, in metagenomes zooplankton can be examined in conjunction with bacteria, archaea, viruses, and protists. Thus, a general decrease in the zooplankton DNA with depth can be seen at HOT, and an increase can be seen in the ODZs (Fig 3 and 5). With amplicons, the data is in % of zooplankton, so these trends in abundance cannot be observed. Additionally, with metagenomes depth profiles of zooplankton and specific bacteria can be compared. However, eDNA amplicons can examine more specific taxonomic levels than we can here at this time [90].

We find significant zooplankton DNA in metagenomic depth profiles, as measured using the phylogenetic placement of reads on a COI gene tree (Fig 1). In both the oxic ocean and ODZs the main zooplankton constituents in metagenomes were crustaceans, Cnidarians, and Gastropods (Fig 3). We were surprised by the abundance of Cnidarians in ODZ metagenomes as they were not mentioned in previous net-based literature [23, 25]. Hydrozoa were the dominant Cnidaria in the ODZ (Fig 3). Hydrozoa have settled hydroid phases that attach to the bottom alternating with very small free-swimming sexually reproducing medusae [91]. We found <5 mm long Cnidaria in UVP images from the ETNP St P2 ODZ (Fig S13). The presence of Cnidaria including Hydrozoa was confirmed in eukaryotic transcriptomes from the ETNP ODZ (Fig 5). Additionally, the few water eDNA amplicon surveys have taken place in the oxic mesopelagic and have also found an abundance of Hydrozoa [90, 92]. Altogether this data indicates that gelatinous organisms should not be ignored when constructing zooplankton-mediated biogeochemical budgets of the ODZ.

Cnidaria have significantly more reads than Crustaceans and Gastropods in metagenomes of all three ODZs (Fig 3). Gelatinous organisms also dominated eRNA transcripts (Fig 4). However, shedding rates of eDNA can vary between species. Jellyfish have a high shedding rate, likely because of their mucus, and crustaceans have a low shedding rate because of their hard carapaces [93]. Thus, the difference in the number of normalized reads seen between Cnidaria and crustaceans in our metagenomic dataset may not reflect an actual over-abundance of Cnidaria. While not indicating organism counts, the amount of zooplankton DNA does indicate the amount of shed organic matter available to heterotrophic bacteria from these sources. The presence of eRNA for a taxa may indicate that cells are fresh and a preferrable food source for bacteria. The fact that we can detect various Cnidaria and copepods in eukaryotic meta-transcriptomes indicates that this zooplankton material is fresh and includes cells that are still active. Though greatly reduced in the ODZ, active cells were still present. Cells shed from zooplankton are likely a food source for heterotrophic bacteria throughout the ocean.

Interestingly, the number of zooplankton reads in metagenomes increased in the ODZs, but the amount of zooplankton reads in transcriptomes decreased (Fig 4). Non-gelatinous zooplankton biomass has previously been shown to decrease in the ETNP ODZ [23]. This could indicate greater preservation of eDNA in the anoxic conditions in the ODZ. Potentially relatedly, there is reduced attenuation of particulate organic matter in ODZs [94], though amino acids are preferentially consumed in the particles [95]. One interpretation of this reduced attenuation is reduced microbial activity degrading complex organic material under anoxic conditions [94]. Thus, anoxia may amplify eDNA signals.

Though we used suspended particulate organic matter to indicate where zooplankton were important sources of organic matter in the ODZ, the bacterial group that we find highly correlated with zooplankton, is SAR11 (Table 1; Fig 5), a free-living group of heterotrophic bacteria that consume simple dissolved organic matter [45]. SAR11 are unlikely to be consuming particulate matter left from zooplankton. While initially surprising, the correlation between SAR11 and zooplankton makes sense as SAR11 bacteria consume small dissolved organic compounds and migrating zooplankton excrete small dissolved organic compounds at depth [35, 36, 45]. Thus, zooplankton at their migration depth are likely providing C and N to microbes in several ways at the same time. In the ETNP, there were quite high R^2^ values between SAR11 and total zooplankton normalized reads in the ODZ (∼0.96 (p=0.0005); Table 1). In the ETSP, St 17 also had high correlations between SAR11 and zooplankton normalized reads (∼0.88 (p=0.003); Table 1), but at St 9 the correlations were still significant (p=0.00002) but R^2^ values were lower (∼0.73; Table 1). In particular, correlations between SAR11 and crustaceans were not significant at ETSP St 9 (Table 1).

We think that this difference between ETSP stations could be due to the quality of organic matter. In the ODZ at St 9 the organic C:N ratios (Fig S9) were similar to algae with plenty of C and N available. However, at St 17, C:N ratios were quite high (10-11) in the ODZ (Fig S9), indicating degraded organic matter with a lack of N. High C:N ratios of organic matter at P2 in the ETNP and St 17 in the ETSP (Fig 2) indicate that at these stations, the SAR11 may have utilized zooplankton excreta as an N source. Ammonia is <10 nM and quite limiting in the offshore ETNP ODZ [51]. Ammonia concentrations were below detection (0.06 µM) in the ETSP ODZ at both St 9 and St 17 (Fig S8) [55]. Both taurine and urea are abundant in zooplankton excreta, though Cnidaria do not excrete urea [34, 35, 39]. Some SAR11 bacteria in the ODZ can utilize urea as an alternative reduced N source, as indicated by the presence of the urease gene [51] (Figure 6). Taurine can also be used as a reduced N source [50]. Active taurine utilization by SAR11 bacteria has been found in surface waters at HOT [49] and across depth profiles in the North Atlantic [48].

Urease genes were present in SAR11 in the ODZs, forming a maximum in each ODZ (Fig 6). These urease maxima are in contrast to HOT where SAR11 urease genes were not found at depth (Fig 6). Ammonia is undetectable in these ODZs, so bacteria may use alternative reduced N sources [51, 55]. The presence of urease genes in the ODZs could be due to the presence of crustacean and gastropod zooplankton in the ODZ, as they excrete urea. However, only 10% of SAR11 had the urease gene in the ETNP ODZ and 5% in the ETSP ODZs (Fig 6). Therefore, 90% of SAR11 did not possess urease genes. Correlations between Cnidaria and SAR11 were particularly high in the ODZs (Table 1). Cnidaria do not excrete urea [39]. Thus, the urease data is consistent with the importance of Cnidaria as a source of organic matter in the ODZ. The highest proportion of SAR11 bacteria with urease genes were actually in surface waters at HOT (Fig 6), where reduced N-containing nutrients can be scarce [84].

Contrastingly, taurine can be considered a direct link between migrating zooplankton and SAR11 because taurine transporter (*tauA*) and C and N assimilation (*tpa*) genes were found in all SAR11 at all stations. The results for *tauD*, representing the pathway to use taurine for organic S assimilation, are more complicated. In the deep oxic waters at HOT and the hypoxic waters below the ETSP St 17 ODZ, the % of SAR11 with *tauD* was elevated, indicating the importance of taurine as an organic S source in those areas. Interestingly the % of SAR11 with the organic S utilization gene *tauD* was reduced both in the ODZs and in surface waters at HOT. Both these areas of the ocean should have additional sources of organic S. Eukaryotic phytoplankton in the surface ocean produce organic sulfonate compounds DMSP, DHSP, cysteate, and isethionate in addition to taurine [49] while sulfate reduction [96, 97] and organic sulfurization occurs in particles in ODZs, leading to the production of sulfonates [97]. Thus, according to gene abundance in the ODZs, taurine was more likely an N and C and energy source than an organic S source for SAR11 there.

Though metagenomes at HOT did not include the zooplankton vertical migration depths (300-500m; [28]), SAR11 and total zooplankton normalized reads still had significant R^2^ values at all three seasons (0.89, 0.85, 0.59; Table 1). Surprisingly, while SAR11 bacteria correlated with zooplankton DNA in HOT metagenomes, they did not correlate with cyanobacteria and only correlated with eukaryotic algae in May (HOT 272) (Table S5). SAR11 bacteria have a streamlined genome and need many exogenous compounds [46]. Scientists have cultured eukaryotic algae and SAR11 bacteria together, and algae provided the SAR11 with their needed S-containing compounds and vitamins [49, 98]. However, in similar cultures, *Prochlorococcus* bacteria were unable to provide SAR11 bacteria with their needed S compounds [99]. We note that HOT is heavily dominated by *Prochlorococcus* [84]. Thus, correlations between SAR11 and eukaryotic algae need to be further investigated in other places where algae are more abundant. We do note that zooplankton eat algae and thus likely excrete some similar compounds as created by algae.

## Conclusions

Zooplankton migrate into ODZs to avoid predators. These zooplankton can bring organic matter and reduced N to the microbial community in these ODZs. We show that zooplankton migration affects suspended particulate organic matter C:N ratios by addition of N. SAR11 bacteria abundance is highly correlated with amounts of zooplankton DNA in metagenomic samples from ODZs and the oxic HOT station. Correlations were particularly high with Cnidarian DNA. We examined SAR11 genes needed to utilize two compounds known to be excreted by zooplankton, taurine, and urea. Cnidaria do not excrete urea but do excrete taurine [35, 39]. Urease was present at 5-10% of the SAR11 population in the ODZs, but was minimal at a similar depth at the oxic HOT station. The presence of urease in SAR11 in the ODZ highlights the importance of excreta of non-Cnidarian zooplankton as a food source there. However, the majority of SAR11 did not possess urease, which could reflect the importance of Cnidaria as a food source. The ability to use taurine as a source of C and N and energy was found in all SAR11, but fewer SAR11 could use taurine as an organic S source in the ODZs, probably because other organic S sources were available. Taurine can be considered a direct link between migrating zooplankton and SAR11 bacteria in ODZs.

## Acknowledgments

This work was funded by Horn Point Laboratory and the University of South Carolina start-up accounts for CAF and XP. We thank Catherine Fitzgerald for filtering eDNA qPCR samples, Hilary Palevsky for creating ETSP satellite chlorophyll maps, Jacob Cram for looking for jellyfish in the UVP images, and Allison Smith for help with 2012 ADCP backscattering data. eDNA qPCR work was funded by the Deerbrook Charitable Trust award DCT 19-34 to LVP. We thank Emily Spady for sampling and prepping ETSP 2013 POM and Jacquelyn Neibauer for sampling 2012 ETNP POM, analyses which were funded by OCE-1153935 to RGK. We thank the captains and crew of the R/V *Revelle*, R/V *Kilo Moana*, R/V *Nathaniel B Palmer*, RV *Sikuliaq*, RV *Thompson* and RV *Sally Ride*. We thank chief scientists Bess Ward, Allan Devol, and Gabrielle Rocap. These cruises were funded by NSF grants OCE-1657663, OCE-1046017, and DEB-1542240 to Bess Ward, Allan Devol, Gabrielle Rocap and Rick Keil.

## Data Availability

Previously published metagenomes can be found at Bioprojects PRJNA704804 (ETSP), PRJNA352737 (HOT), and PRJNA350692 (ETNP). Assembled ETSP contigs have been added to Bioproject PRJNA704804 and are accession numbers JAMYFK000000000-JAMYGN000000000. ETNP eukaryotic transcript reads can be accessed from NCBI SRA accession numbers SAMN30032067 to SAMN30032098.

