## Supplementary Materials for "Linking mesozooplankton and SAR11 bacteria in Oxygen Deficient Zones and the open ocean"

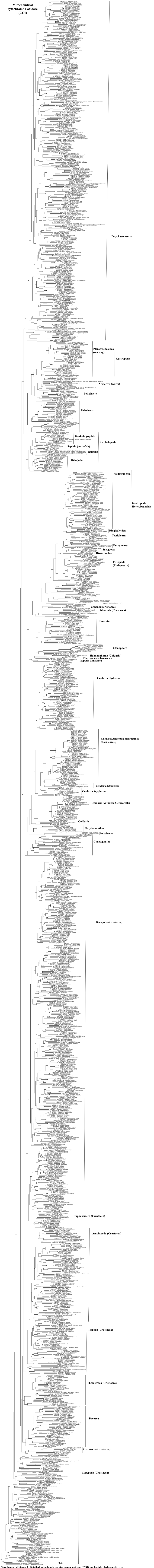


**Supplemental Figure 1. Detailed mitochondria cytochrome oxidase (COI) nucleotide phylogenetic tree.**

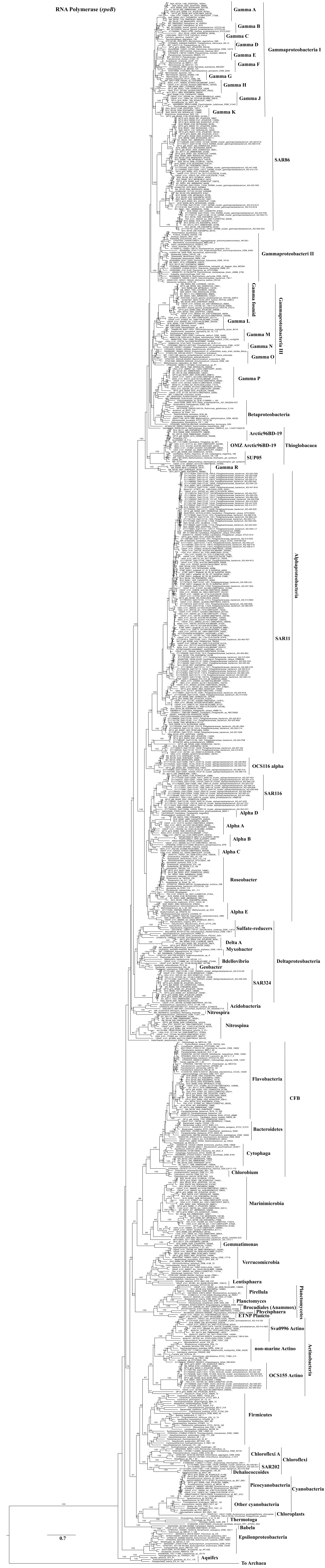


**Supplemental Figure 2. RNA polymerase (*rpoB*) amino acid phylogenetic tree. Numbers on the nodes indicate bootstrap values.**

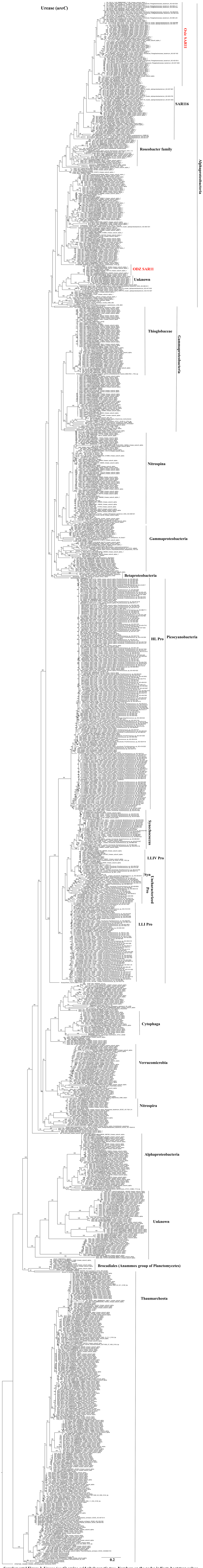


**Supplemental Figure 3. Urease (*ureC*) amino acid phylogenetic tree. Numbers on the nodes indicate bootstrap values.**

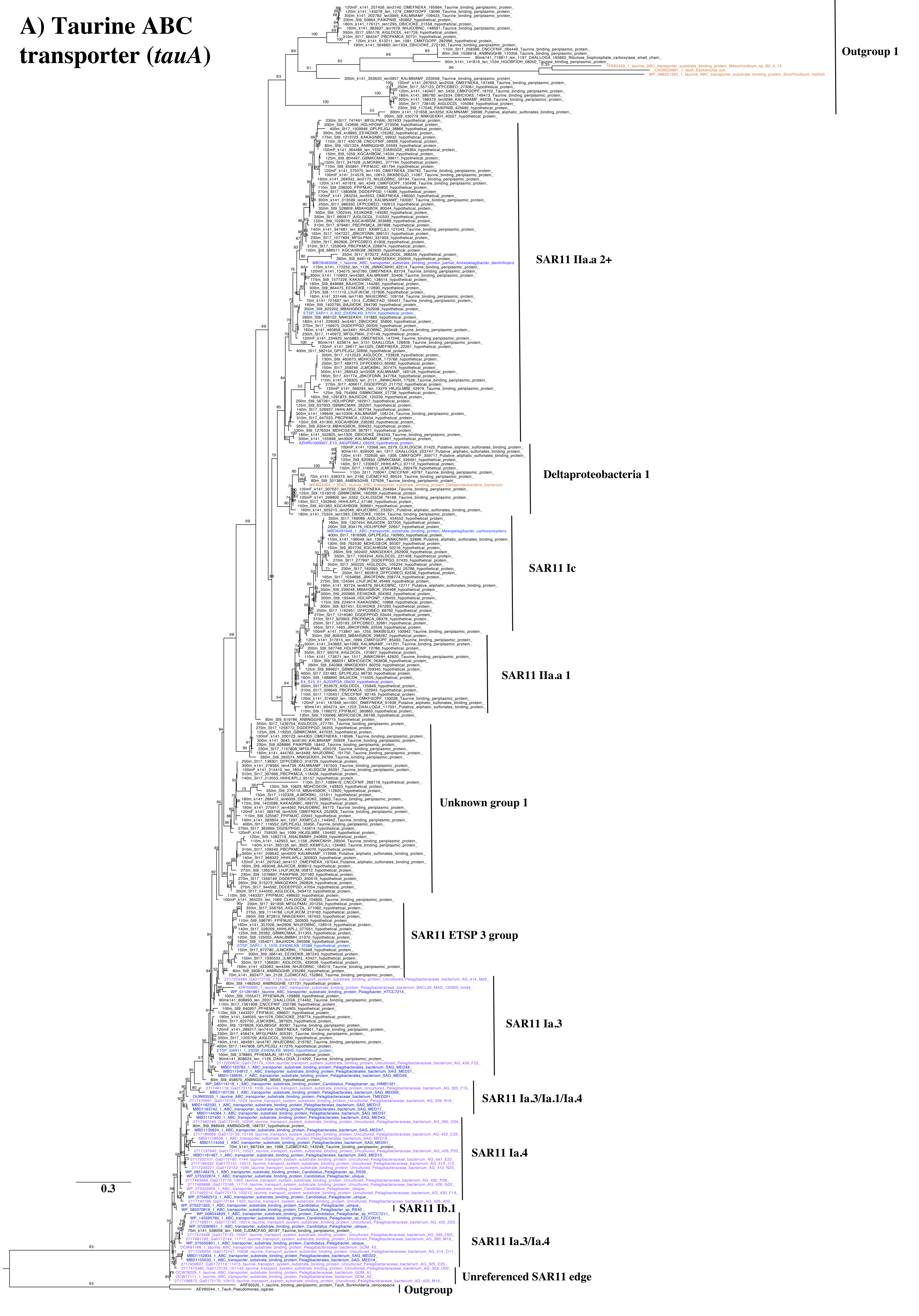
A) Taurine ABC transporter (tauA)
Outgroup 1
SAR11 IIa.a 2+
Deltaproteobacteria 1
SAR11 Ic
SAR11 IIa.a 1
Unknown group 1
SAR11 ETSP 3 group
SAR11 Ia.3
SAR11 Ia.3/Ia.1/Ia.4
SAR11 Ia.4
SAR11 Ib.1
SAR11 Ia.3/Ia.4
Unreferenced SAR11 edge
Outgroup
0.3

### B) Taurine dioxygenase (*tauD*)

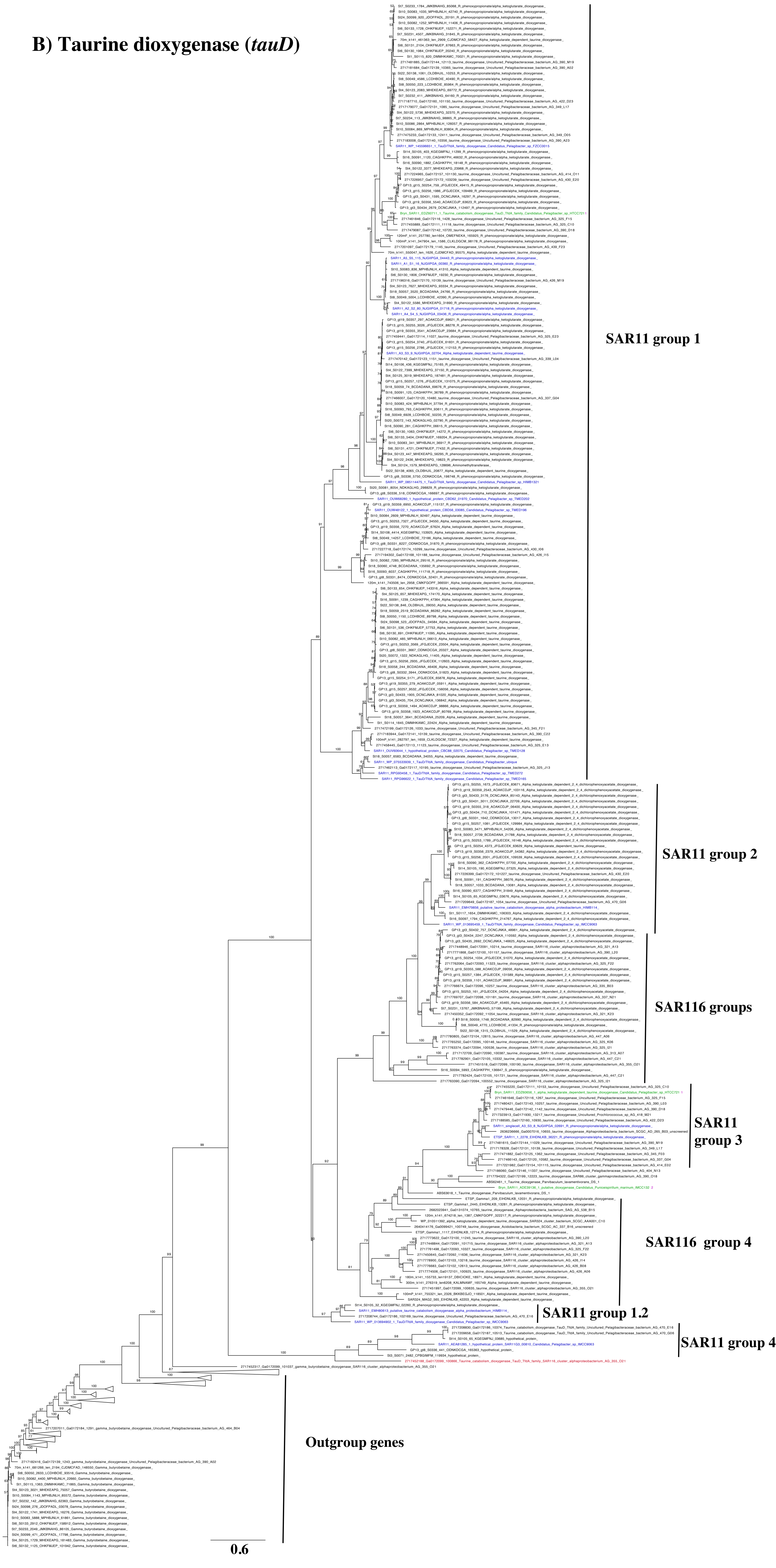

**C) tpa**

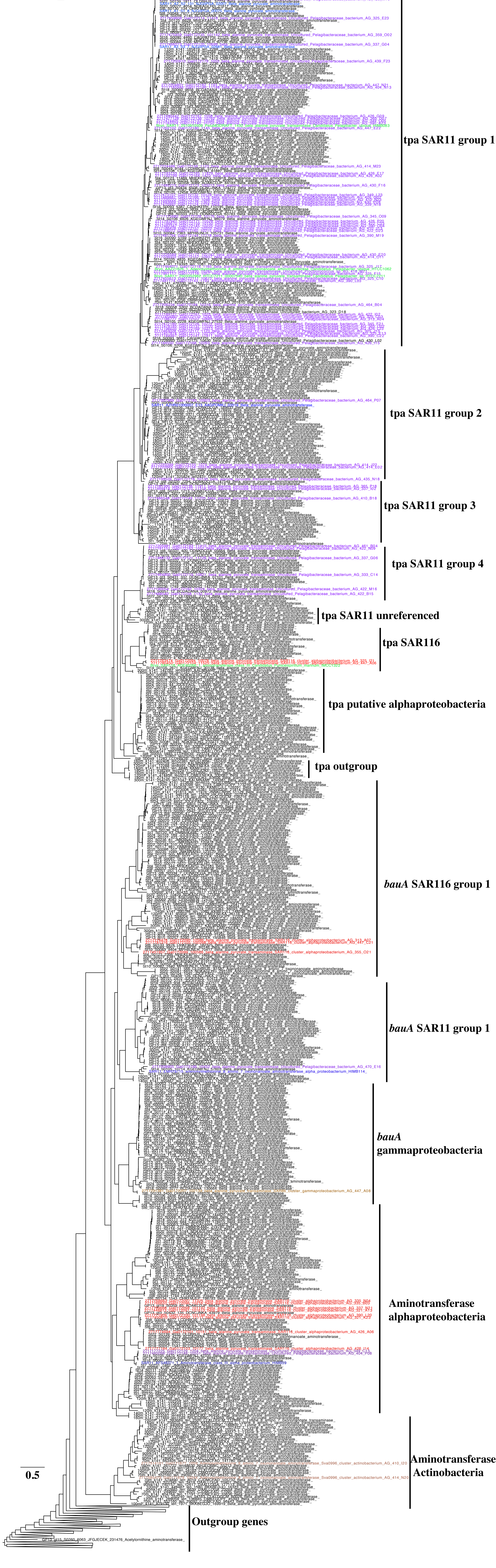

tpa SAR11 group 1
tpa SAR11 group 2
tpa SAR11 group 3
tpa SAR11 group 4
tpa SAR11 unreferenced
tpa SAR116
tpa putative alphaproteobacteria
tpa outgroup
*bauA* SAR116 group 1
*bauA* SAR11 group 1
*bauA* gammaproteobacteria
Aminotransferase alphaproteobacteria
Aminotransferase Actinobacteria
Outgroup genes
0.5

Supplemental Figure 4. Taurine metabolism amino acid phylogenetic trees for A) substrate binding subunit of ABC transporter *tauA*, B) organic S assimilation gene *tauD*, and C) energy providing C and N assimilation gene *tpa*. Blue and purple indicate SAR11 references. Red indicates SAR116 references. Green indicates references from (Durham *et al.*, 2019) used to check gene identity. Brown indicates other references. Numbers on the nodes indicate bootstraps.

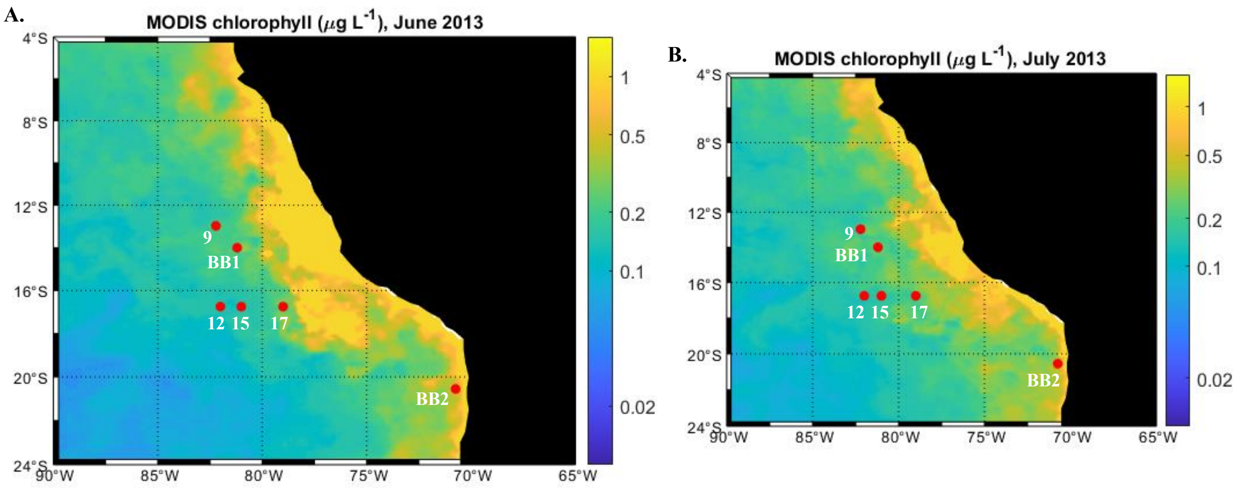


| **C.** | St 9 | BB1 | St 12 | St 15 | St 17 | BB2 |
|---|---|---|---|---|---|---|
| Latitude N | -13.00 | -14.00 | -16.74 | -16.74 | -16.75 | -20.76 |
| Longitude E | -82.20 | -81.20 | -82.00 | -81.00 | -79.00 | -70.73 |
| Date sampled | 7/7/2013 | 7/1/2013 | 7/10/2013 | 7/10/2013 | 7/11/2013 | 7/15/2013 |
| June Chl a (μg/L) | 0.20 | 0.24 | 0.15 | 0.15 | 0.3 | 0.49 |
| July Chl a (μg/L) | 0.22 | 0.23 | 0.17 | 0.17 | 0.23 | 0.45 |

Supplemental Figure 5. Map of MODIS satellite chlorophyll from A) June 2013 and B) July 2013 with ETSP stations (red dots). C) Surface satellite chlorophyll from each station and the date sampled. Map Credit: Hilary Palevsky.

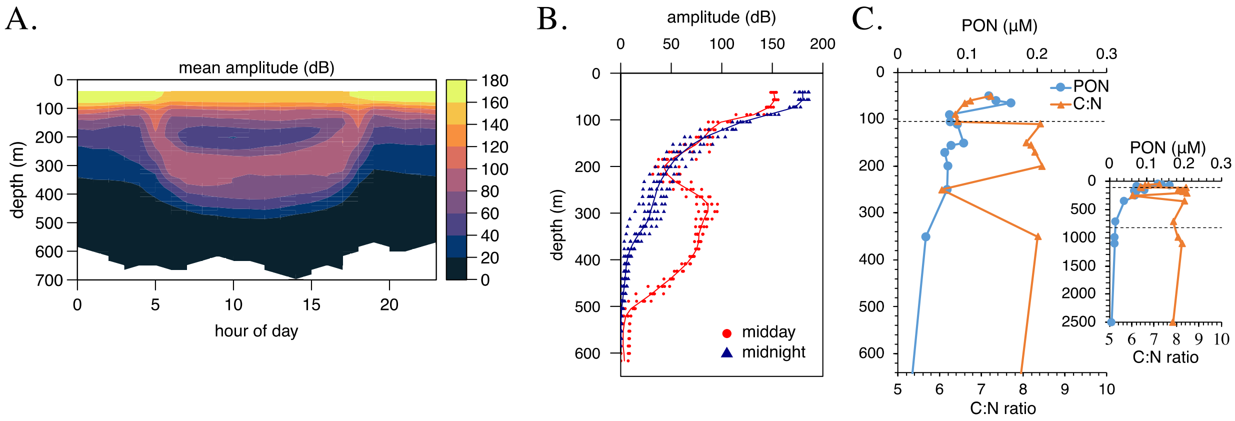


Supplemental Figure 6. ETNP St P2 in April 2012 A) Mean amplitude of ADCP backscattering over a 24 hour period, indicating zooplankton migration, B) Comparison of backscattering amplitude at noon and midnight over a 5 day period, C) Suspended particulate organic N concentrations and C:N ratios over the top 600m. Inset: Suspended particulate organic N concentrations and C:N ratios over the entire water column. Dashed line indicates the boundaries of the ODZ.

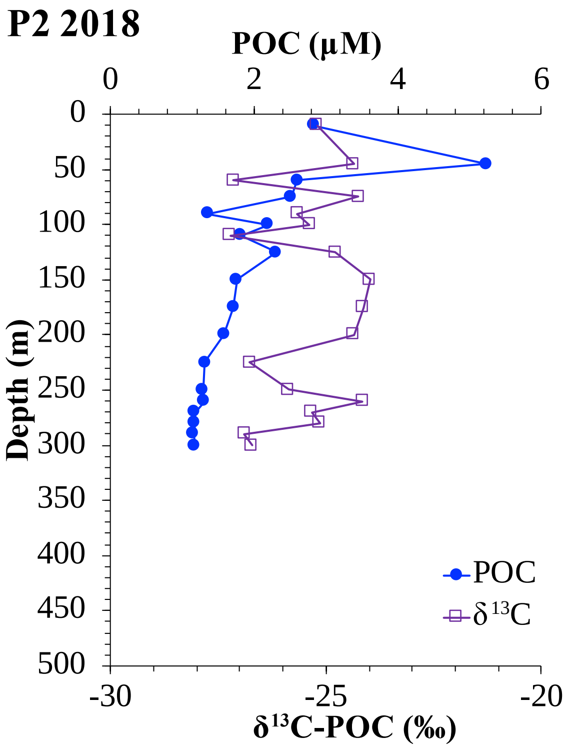


Supplemental Figure 7. Suspended POC concentrations and $\delta^{13}C$ from ETNP St P2 in 2018.

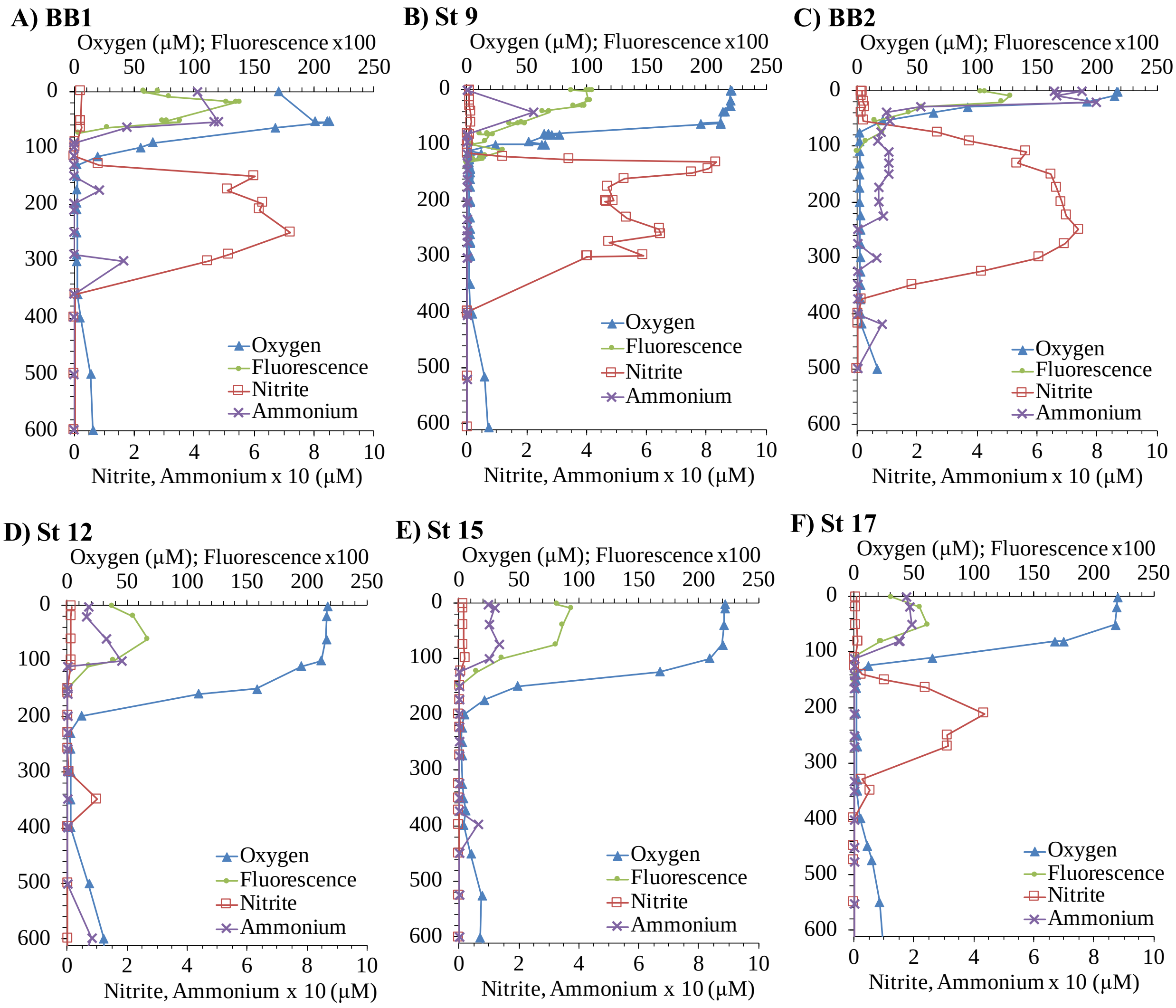


Supplementary Figure 8. Oxygen, nitrite, ammonium and fluorescence depth profiles for stations with POM data. A) St BB1, B) St 9, C) BB2, D), St 12, E) St 15, F) St 17. Oxygen and nitrite data are previously published in (Peters *et al.*, 2018) and ammonium is previously published in (Widner *et al.*, 2018). The detection limit for ammonium was 0.06 µM and values below that limit were set at zero. Ammonium values were multiplied by 10 to better fit on the axis scale. See the map in Figure S5 for station locations.

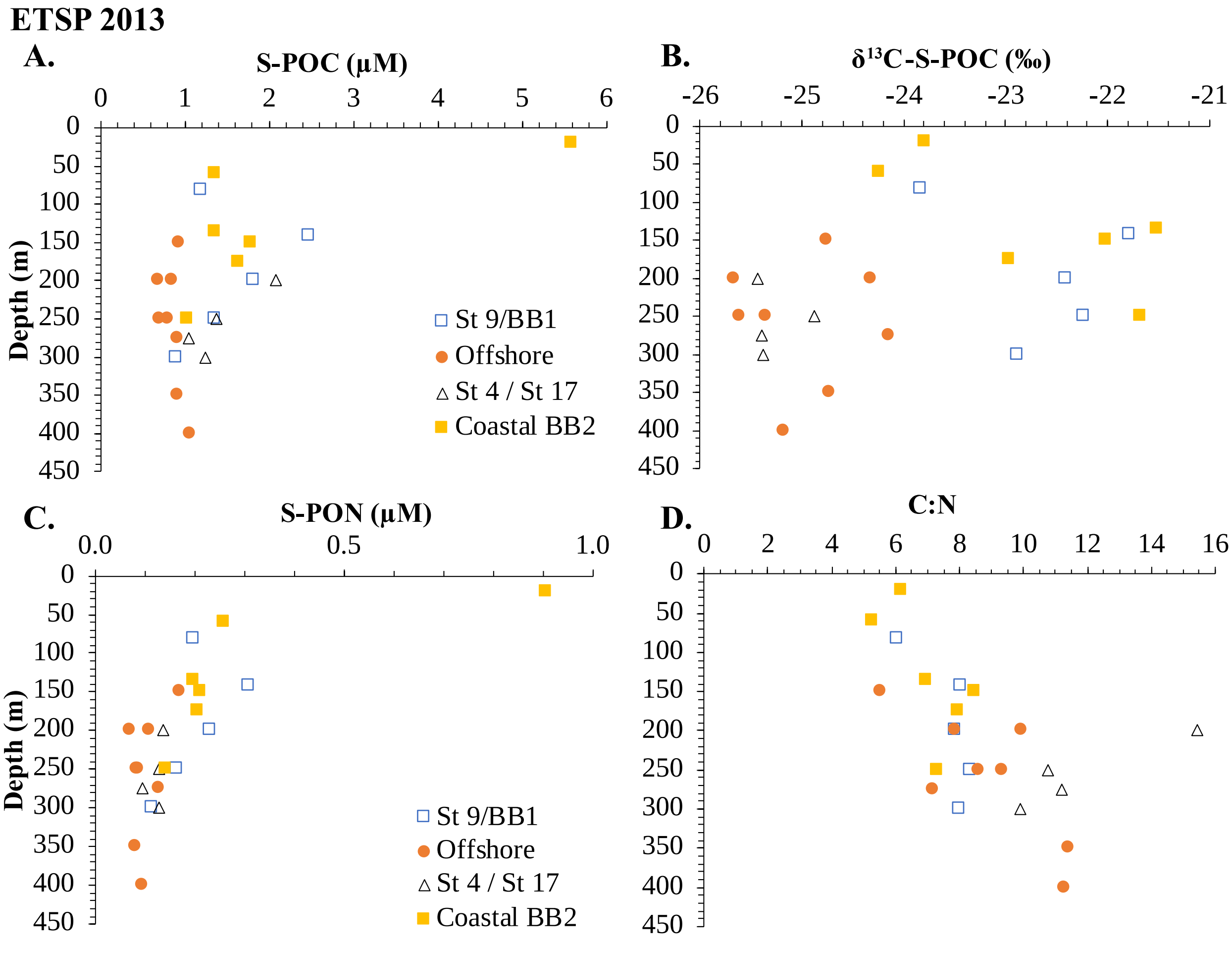


Supplemental Figure 9. ETSP 2013 organic matter (>0.7 µm). A) suspended particular organic C, B) $\delta^{13}C$ of suspended POC, C) suspended particular organic N, D) C/N ratios of suspended POM. St 9 and BB1 were combined. St 12 and 15 are combined and called offshore. BB2 is considered coastal. For more information about these stations see Figure S7 and Peters et al (2018).

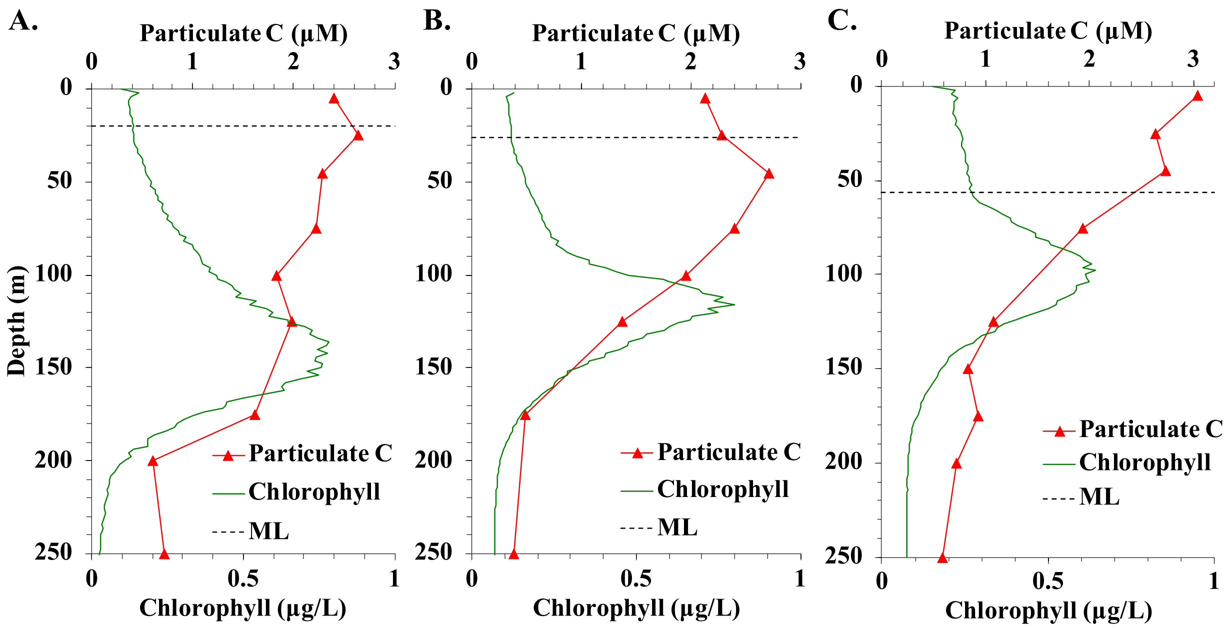


Supplemental Figure 10. Particulate carbon and chlorophyll at A) HOT 272 (May), B) HOT 275 (Aug), and C) HOT 278 (Nov). Data from (Luo *et al.*, 2020). Dotted lines indicate the base of the mixed layer. Particulate carbon includes both organic and inorganic C.

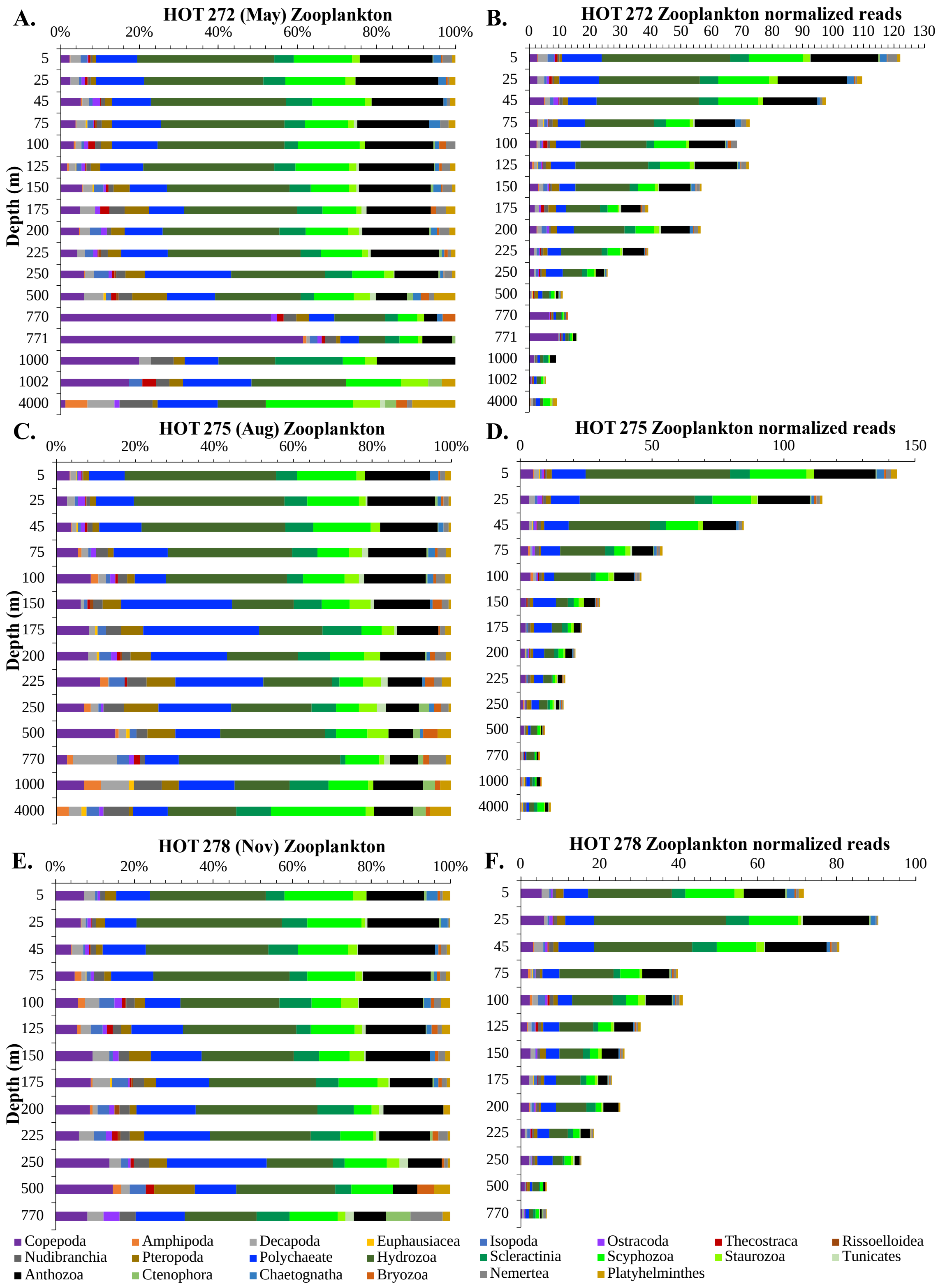


Supplemental Figure 11. Zooplankton groups for the COI gene from metagenomic depth profiles were examined as % of zooplankton community and by normalized reads for A&B) HOT 272, C&D) HOT 275, and E&F) HOT 278.

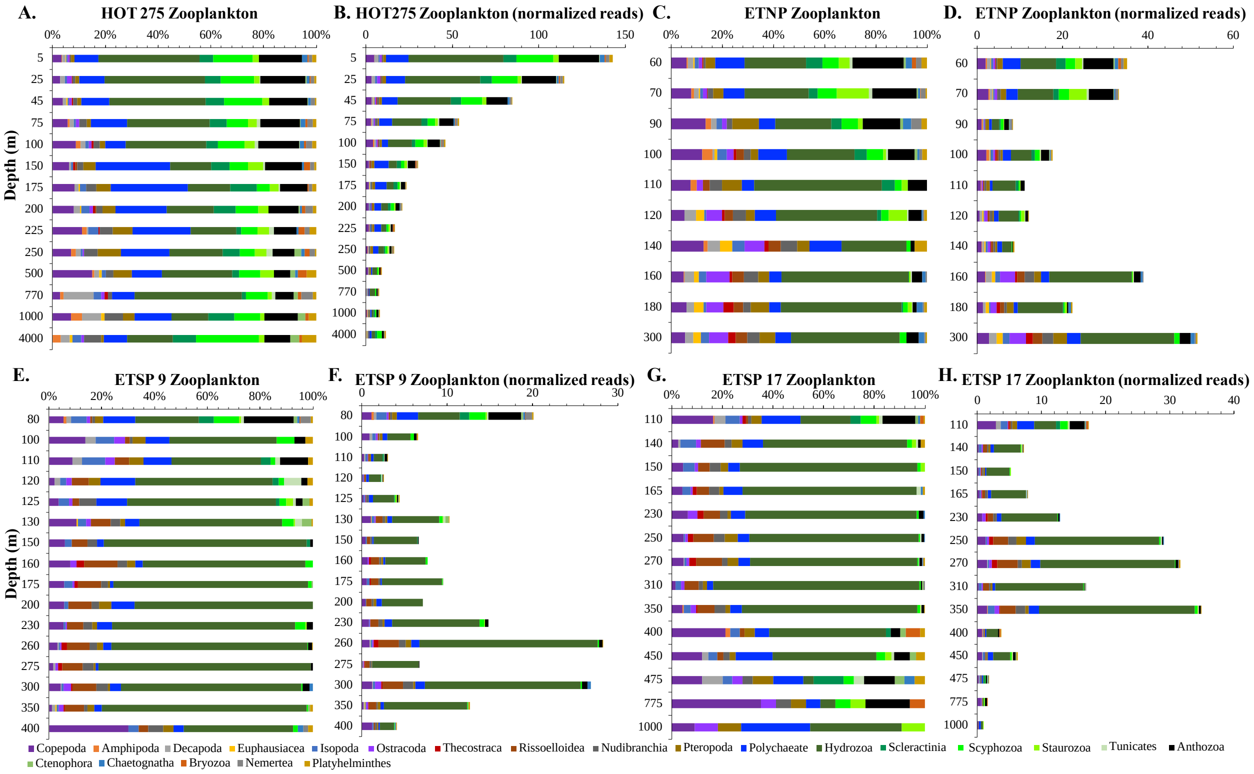


Supplemental Figure 12. Zooplankton groups for the COI gene from metagenomic depth profiles were examined as % of zooplankton community and normalized reads for A&B) HOT 275 (Aug), C&D) ETNP, E&F) ETSP St 9, and G&H) ETSP St 17.

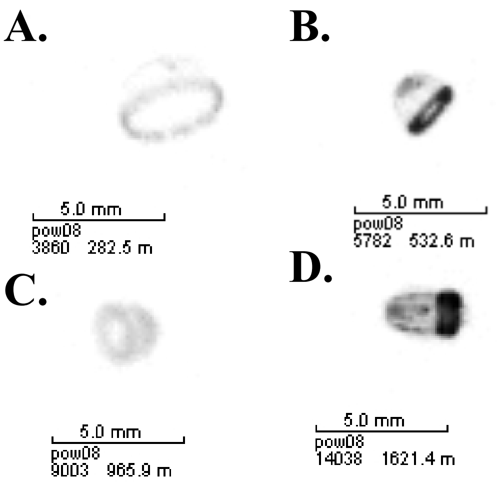


Supplemental Figure 13. UVP images from ETNP St P2 in 2017 (dataset published in (Cram *et al.*, 2022)) at A) 282m in the ODZ, B) 532m in the ODZ, C) 965m in oxic waters below the ODZ, and D) 1621m in oxic waters below the ODZ. The ruler indicates 5 mm in length.

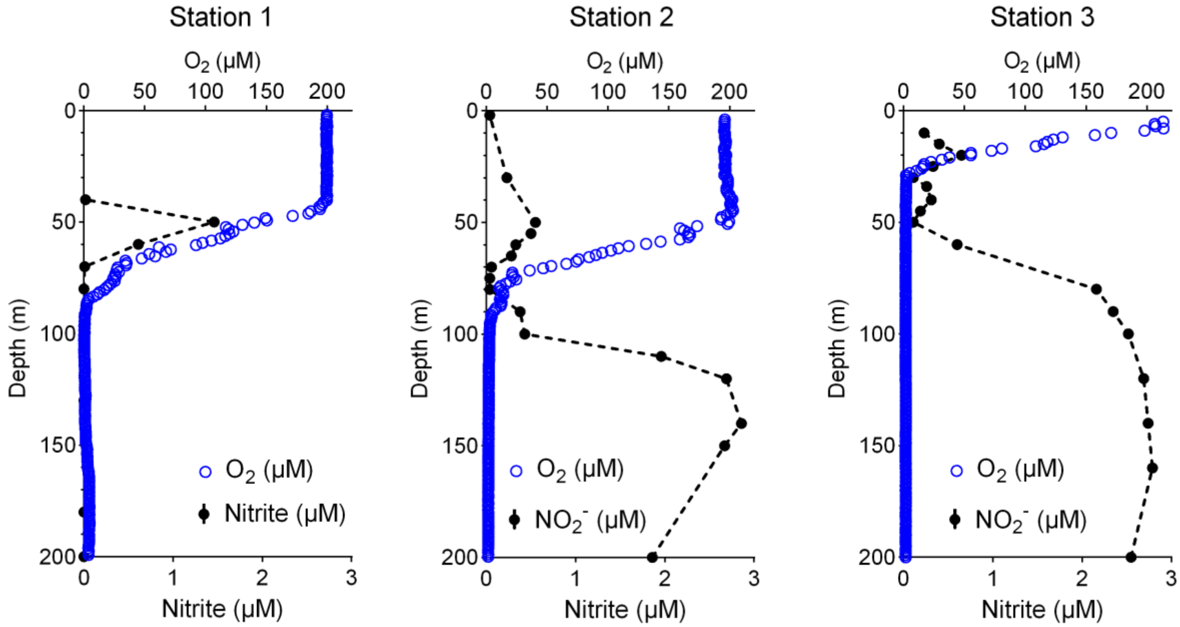


Supplemental Figure 14. Oxygen and nitrite profiles for ETNP Stations 1-3 in 2018 sampled for transcriptomics. Data was previously published in (Kelly *et al.*, 2021).

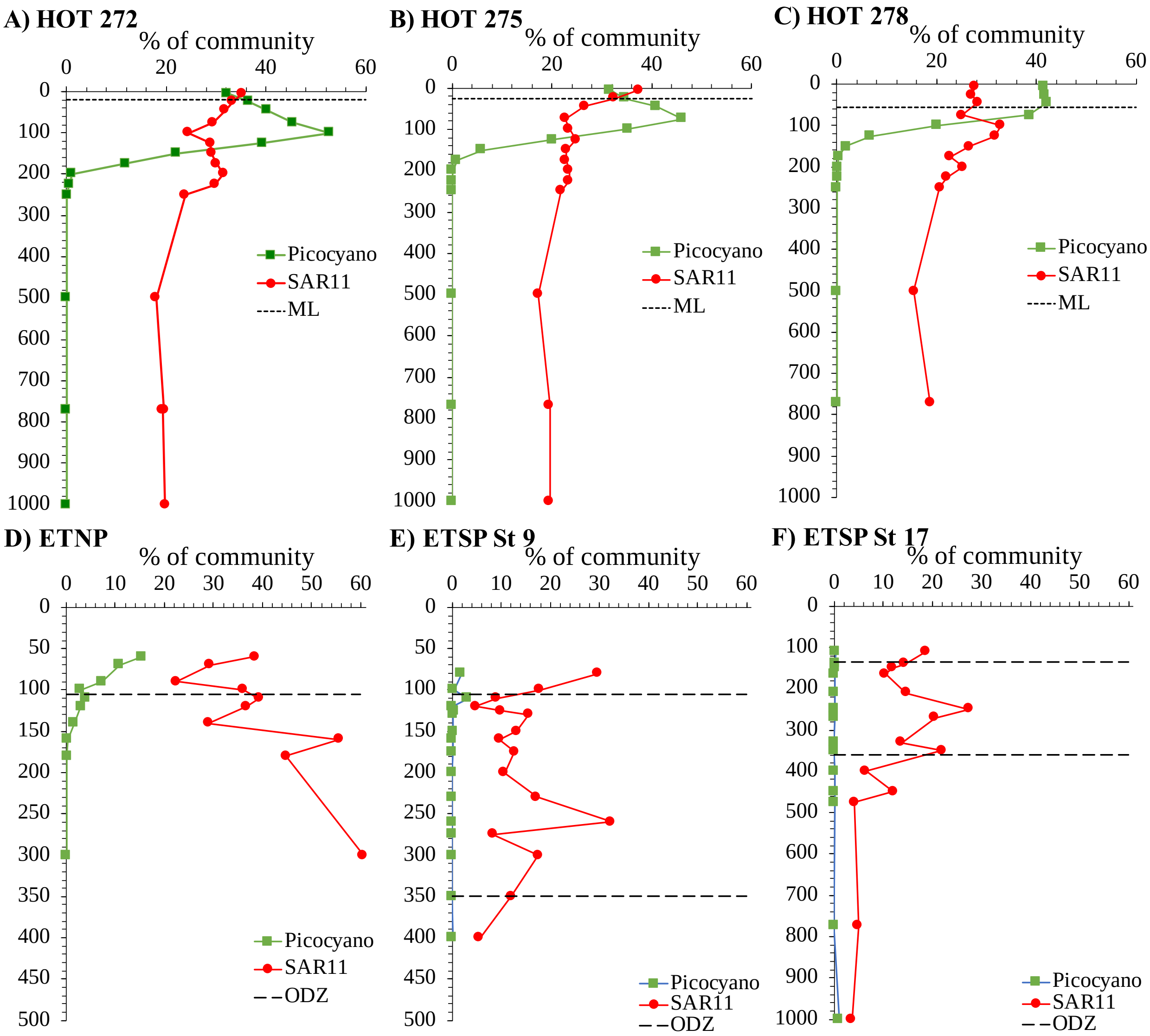


Supplemental Figure 15. SAR11 and Picocyanobacteria (*Prochlorococcus* and *Synechococcus*) % community (*rpoB*) at A) HOT 272, B) HOT 275, C) HOT 278, D) ETNP, E) ETSP St 9, and F) ETSP St 17. Dotted lines indicate the base of the mixed layer. Dashed lines indicate the top and bottom of the ODZ.

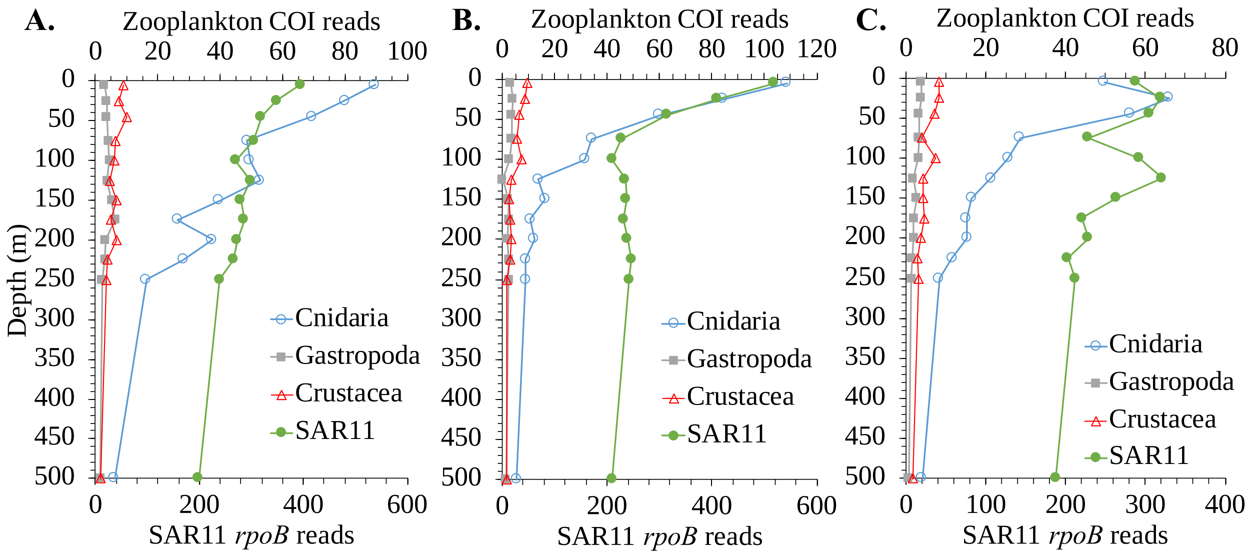


Supplemental Figure 16. Depth profiles of Cnidaria, Gastropoda, and Crustacea (COI) and SAR11 (*rpoB*) metagenomic reads in the A) HOT 272, B) HOT 275, and C) HOT 278.

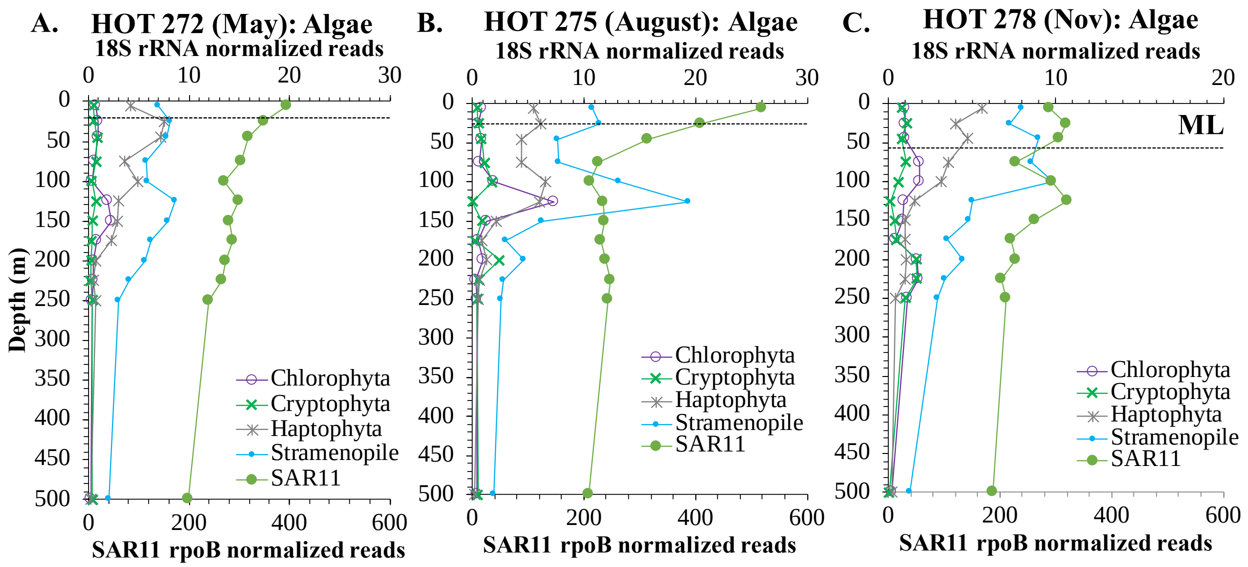


Supplemental Figure 17. Depth profiles of eukaryotic algae (18S rRNA normalized reads) from Fuchsman et al (2022) and SAR11 *rpoB* normalized reads for A) HOT 272, B) HOT 275, and C) HOT 278.

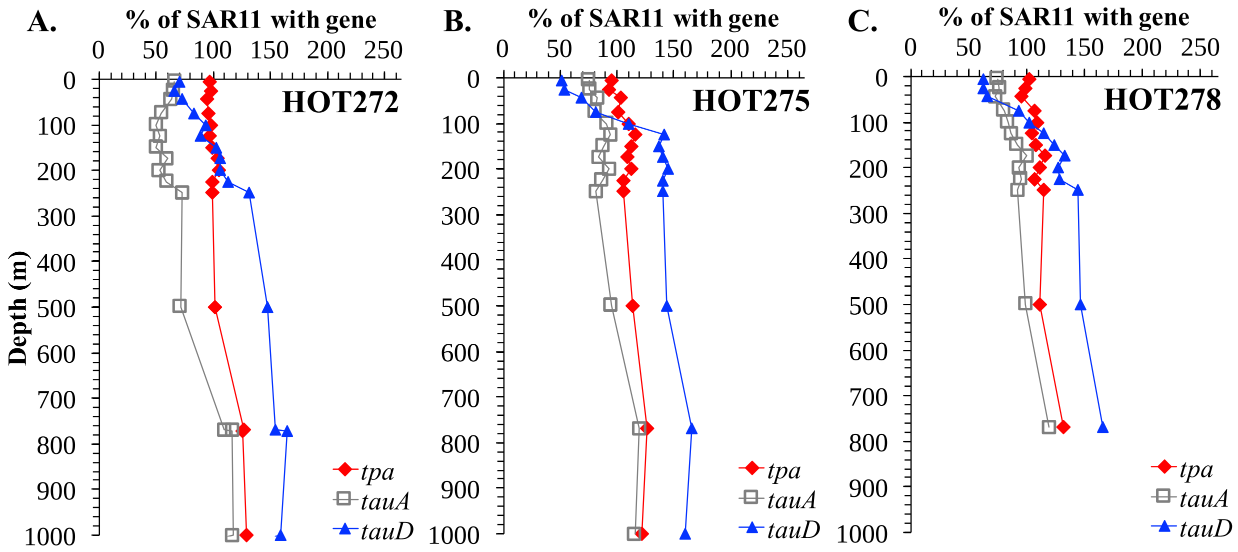


Supplemental Figure 18. Metagenomic depth profiles of taurine metabolism genes taurine: pyruvate transaminase (*tpa*), taurine dioxygenase (*tauD*), taurine ABC transporter substrate-binding subunit (*tauA*) in SAR11 for A) HOT 272 (May) B) HOT 275 (Aug), and C) HOT 278 (Nov).

**Supplemental Table 1.** List of stations and measurements at each station.

| Station | Latitude/Longitude | Meta-genomics | Meta-transcriptomics | POM | qPCR of eDNA |
|---|---|---|---|---|---|
| ETNP St 136 | 17.04 ºN 106.54 ºW | X | | X | |
| ETNP St P2 | 16.9ºN 107ºW | | | X | X |
| ETNP St 1 | 10°N 113°W | | X | | |
| ETNP St 2 | 15.77°N 105°W | | X | | |
| ETNP St 3 | 17.68°N 102.35°W | | X | | |
| ETSP St 9 | 13ºS 82.2ºW | X | | X | |
| ETSP St 17 | 16.7ºS 79ºW | X | | X | |
| HOT | 22º45'N 158ºW | X | | X | |

**Supplementary Table 4.** Number of raw reads in each metatranscriptome library.

| Sample | Station | Cast number | Depth (m) | Number of raw reads |
|---|---|---|---|---|
| Cast6_60m | 1 | 6 | 60 | 15,308,412 |
| Cast18_110m | 1 | 18 | 110 | 14,949,138 |
| Cast19_90m | 1 | 19 | 90 | 14,570,593 |
| Cast20_54m | 1 | 20 | 54 | 16,762,062 |
| Cast22_60m | 1 | 22 | 60 | 15,029,719 |
| Cast23_40m | 1 | 23 | 40 | 16,691,176 |
| Cast27_275m | 2 | 27 | 275 | 15,538,952 |
| Cast31_90m | 2 | 31 | 90 | 12,876,070 |
| Cast34_60m | 2 | 34 | 60 | 15,368,312 |
| Cast35_120m | 2 | 35 | 120 | 12,331,170 |
| Cast37_92m | 2 | 37 | 92 | 10,675,870 |
| Cast39_85m | 2 | 39 | 85 | 10,094,611 |
| Cast41_90m | 2 | 41 | 90 | 13,659,711 |
| Cast47_113m | 2 | 47 | 113 | 12,453,194 |
| Cast48_90m | 2 | 48 | 90 | 13,912,038 |
| Cast50_40m | 2 | 50 | 40 | 13,126,731 |
| Cast52_65m | 2 | 52 | 65 | 14,889,047 |
| Cast55_120m | 2 | 55 | 120 | 16,914,741 |
| Cast59_250m | 2 | 59 | 250 | 12,767,179 |
| Cast59_30m | 2 | 59 | 30 | 13,432,084 |
| Cast59_900m | 2 | 59 | 900 | 12,769,528 |
| Cast63_10m | 2 | 63 | 10 | 13,215,238 |
| Cast72_45m | 3 | 72 | 45 | 12,054,655 |
| Cast73_16m | 3 | 73 | 16 | 11,996,691 |
| Cast76_36m | 3 | 76 | 36 | 12,654,069 |
| Cast80_50m | 3 | 80 | 50 | 13,710,020 |
| Cast82_48m | 3 | 82 | 48 | 15,105,664 |
| Cast85_20m | 3 | 85 | 20 | 12,708,093 |
| Cast93_10m | 3 | 93 | 10 | 14,368,100 |
| Cast97_40m | 3 | 97 | 40 | 11,546,283 |
| Cast98_25m | 3 | 98 | 25 | 13,498,132 |
| Cast101_80m | 3 | 101 | 80 | 13,270,574 |

**Supplemental Table 5. Correlations between SAR11 (*rpoB*) normalized reads and eukaryotic algae 18S rRNA normalized reads (Fuchsman et al 2022) or cyanobacteria (*rpoB*) normalized reads. HOT 278 has 3 points in the mixed layer (56m), which affects the regression. The regression HOT 278 ML has only one data point in the mixed layer. Slope and intercept were not provided for regressions with a p value ≥0.05.**

| | HOT 272 (May) | HOT 275 (Aug) | HOT 278 (Nov) | HOT 278 ML |
|---|---|---|---|---|
| **Cyanobacteria** | | | | |
| slope | | | 2±1 | |
| intercept | | | -530±270 | |
| $R^2$ | 0.33 | 0.34 | 0.41 | 0.24 |
| P value | 0.05 | 0.06 | 0.02 | 0.15 |
| **Total Eukaryotic Phytoplankton** | | | | |
| slope | 0.07±0.02 | | 0.06±0.02 | |
| intercept | -10±5 | | -6±6 | |
| $R^2$ | 0.55 | 0.2 | 0.40 | 0.32 |
| P value | 0.006 | 0.2 | 0.03 | 0.08 |
| **Stramenopile** | | | | |
| slope | 0.030±0.009 | | 0.04±0.01 | 0.04±0.02 |
| intercept | -2.8±2.6 | | -4±3 | -5±4 |
| $R^2$ | 0.53 | 0.07 | 0.49 | 0.44 |
| P value | 0.007 | 0.1 | 0.01 | 0.03 |
| **Haptophyta** | | | | |
| slope | 0.03±0.01 | | 0.025±0.009 | |
| intercept | -7±3 | | -4±2 | |
| $R^2$ | 0.48 | 0.29 | 0.44 | 0.36 |
| P value | 0.01 | 0.08 | 0.02 | 0.06 |
| **PON** | | | | |
| slope | 0.0017±0.0006 | | | |
| intercept | -0.2±0.1 | | | |
| $R^2$ | 0.53 | 0.11 | 0.41 | 0.24 |
| P value | 0.03 | 0.4 | 0.06 | 0.2 |
| **POC** | | | | |
| slope | 0.012±0.003 | | | |
| intercept | -2±1 | | | |
| $R^2$ | 0.61 | 0.17 | 0.36 | 0.22 |
| P value | 0.01 | 0.3 | 0.08 | 0.3 |
